# The effect of grape juice dilution on oenological fermentation

**DOI:** 10.1101/2020.07.29.226142

**Authors:** Jennifer Margaret Gardner, Michelle Elisabeth Walker, Paul Kenneth Boss, Vladimir Jiranek

**Affiliations:** Department of Wine and Food Science, School of Agriculture, Food and Wine, The University of Adelaide, Waite Campus, Urrbrae, SA 5064, Australia; CSIRO, Agriculture and Food, Waite Campus, Urrbrae, SA 5064, Australia; Australian Research Council Training Centre for Innovative Wine Production, The University of Adelaide, Waite Campus, Urrbrae, SA 5064, Australia

**Keywords:** fermentation, dilution, yeast, nutrient, volatile compounds

## Abstract

The impact of water addition to grape juice in winemaking, on both alcoholic and malolactic fermentation duration and outcome has been examined using commercial wine yeasts, Lalvin EC1118™ and Lalvin R2™ and malolactic bacteria Lalvin VP41™. As expected, dilution with water did not impede fermentation, instead resulted in shortened duration, or in the case of malolactic fermentation enabled completion in these conditions. Addition of complex organic nutrient further shortened alcoholic fermentation by Lalvin R2™ and in some conditions also reduced the duration of malolactic fermentation. In general, volatile compounds and some major yeast metabolites were present at lower concentrations at the end of fermentation where juices were diluted and the addition of organic complex nutrient also influenced the concentration of some compounds in wine. These findings are significant to commercial winemaking, highlighting that winemakers should consider potential impacts of juice dilution on processing efficiencies along with wine flavour and aroma.

**Highlights: Gardner et al. The effect of grape juice dilution on fermentation:**

- Grape juice dilution shortened both alcoholic and malolactic fermentation
- In some conditions addition of commercial nutrient decreased fermentation duration
- In general wine volatiles decrease with grape juice dilution
- Isoamyl acetate can be decreased in wine by grape juice dilution

## 1. Introduction

The addition of water pre-fermentation is legal and commonplace in many countries, including more recently, Australia. The wording of legislature around water additions varies between countries, for instance additions are allowed “to facilitate fermentation” (USA, Australia), “ pre-fermentation and not to below 13.5 °Bé” (Australia) or “where required on account of a specific technical necessity” (European Union). Fermentation difficulties can be encountered in high sugar musts where yeasts can fail to ferment all available sugars, leading to wines that are out of specification. High-sugar musts are becoming increasingly common since elevated daily average temperatures during ripening (Schultz, 2016) can lead to accelerated and uneven phenological development of grapes, resulting in vintage compaction and delay of harvest by winemakers waiting for “flavour ripeness”. Market limitations also exist for higher alcohol wines, where higher taxes are incurred and some consumers are increasingly avoiding these in favour of wines with lower alcohol, suiting some modern flavour and healthier lifestyle choices. The addition of water to must pre-fermentation to combat the problems associated with high sugar musts is a simple and inexpensive procedure for which the practical logistics have been discussed (Cowey, 2017). Previous studies have examined the effect of juice dilution with water on final concentrations and sensory impacts of various compounds in wine (Harbertson, Mireles, Harwood, Weller, & Ross, 2009; Petrie, Teng, Smith, & Bindon, 2019; Schelezki, Antalick, Šuklje, & Jeffery, 2020; Schelezki, Smith, Hranilovic, Bindon, & Jeffery, 2018; Schelezki, Suklje, Boss, & Jeffery, 2018; Teng, Petrie, Smith, & Bindon, 2020). Authors report variable effects on wine, from an overall decrease in wine volatiles (Schelezki et al., 2020) to an increase to specific compounds (Schelezki, Suklje, et al., 2018), and minor to measurable decreases in colour, tannin and phenolics (Schelezki, Smith, et al., 2018; Teng et al., 2020). Some wines were also reported as generally not being reduced in sensorial complexity, in fact some dilutions maintain many of the fuller bodied and richer flavours of wine from undiluted juices (Petrie et al., 2019). In terms of grape derived compounds, presumably this reflects that changing the juice to solid ratio, or indeed simply providing more water, enables increased extraction. The variability in the reported effect of dilution inevitably arises due the complexities of variety, vintage and experimental differences, as reported by (Schelezki et al., 2020). Overall, authors suggest that impacts on wine of juice dilution pre-fermentation are surprisingly minor. Similarly it is hypothesised that these additions will not impede alcoholic and malolactic fermentation, however research has yet to specifically address this. We hypothesise that the addition of water to juice pre-fermentation will shorten fermentation time and effect the final concentration of some compounds in the final wine in proportion to the dilution. To test this, we analysed fermentation dynamics of two commercial wine yeast; Lalvin EC1118™ (aromatically neutral) and Lalvin R2™ (aromatic) over a range of juice dilutions (16 Bé juice diluted to 14.5, 13.5 and 12.5 Bé, corresponding to water additions of 10.5, 17.2 and 24%). Pressed juice from white grapes was chosen so as to avoid the complexities introduced when winemaking with skin contact is undertaken. Furthermore, studies addressing the impact of juice dilution on extraction of colour and phenolics in red wines fermented in the presence of skins have been recently reported (Schelezki et al., 2020; Teng et al., 2020). Nitrogen was ameliorated to be equal across all juices to ensure this was not impacting fermentation dynamics. Yeast nutrient (FERMAID^®^ O, an organic complex nutrient; Lallemand) was also added to examine if any negative effects (potentially due to the dilution of micronutrients), could be reduced. It is well known that many compounds (volatile and non-volatile) in wine change with the addition of nitrogenous compounds (Torrea, Fraile, Garde, & Ancin, 2003; Bell & Henschke, 2005), thus inclusion of this analysis allowed us to examine the combined effects of dilution and nutrient supplementation.

## 2. Materials and methods

### 2.1 Strains and media

*Saccharomyces cerevisiae* strains Lalvin EC1118™ and Lalvin R2™ (Lallemand, Canada) were chosen due to their common commercial use, variation in nutrient requirements and contribution to wine aroma (Lavin EC1118™ regarded as neutral and Lalvin R2™ as aromatic). Yeast were grown from single colonies in Yeast Peptone Dextrose (YPD) medium (1% w/v yeast extract, 2% w/v bacteriological peptone, and 2% w/v glucose) overnight at 28 °C with shaking (120 rpm). These cells were then inoculated at 2.5 x 10^6^ cells mL^−1^ in diluted juice (45%, sterile (0.22 μm), 2018 Viognier-Marsanne blend, 10% YPD, 45% water) and grown overnight. This culture was used to inoculate experimental fermentations.

*Oenococcus oeni* malolactic bacteria strain VP41™ (Lallemand) was chosen as it is widely used in the Australian wine industry. VP41 (2.5g) was rehydrated from a commercial packet, according to the manufacturer’s instructions and grown in 50 mL of MRSAJ (de Man-Rogosa-Sharp medium (Amyl Media) supplemented with 20% (^v^/_v_) apple juice and 0.1% cyclohexamide (^v^/_v_)) for 4 days at 30 °C and 20% CO_2_. The culture was centrifuged (10 min, 4600 x *g*), and the pellet was washed in phosphate buffered saline (PBS) and re-centrifuged. The cell pellet was washed with 50 mL Viognier-Marsanne juice (13.5 Bé, 0.22 μm), spun down and resuspended in 50 mL Viognier-Marsanne juice. Light microscopy confirmed the culture as viable *O. oeni*. The culture was grown overnight at 30 °C and 20% CO_2_ prior to inoculation at 1:100 (0.25 mL) in the experimental fermentations. Culture viability was analysed using spot plating (10 uL of serially diluted cultures were grown on MRSAJ with 2% agar, and colonies enumerated post growth).

A filter sterile (0.22 μm) 2018 Viognier-Marsanne (~75% Viognier) blended juice was used for experimental fermentations (16 Bé, pH 3.2, 3.13 g L^−1^ malic acid, 18:63 mg L^−1^ free:total SO_2_, 60 mg L^−1^ ammonia, 196 mg L^−1^ alpha amino nitrogen, 245 mg L^−1^ yeast assimilable nitrogen (YAN)). Juices were diluted with ultrapure water (pH 7.0) to 14.5, 13.5 and 12.5 Bé (10.5, 17.2 and 24% water). Density after addition of water was calculated instead of measured as, based on experience, measurement at that stage is inaccurate. Nitrogen was also adjusted in diluted juices to 245 mg YAN L^−1^ with diammonium phosphate and complex organic nutrient (FERMAID^®^ O, Lallemand) was added where indicated at either 0 (control), 200 or 400 mg L^−1^.

### 2.2 Experimental fermentation

Fermentations were conducted with a custom-made high throughput robotic platform ‘Tee-bot v.2.0’ built on an EVO freedom workdeck (Tecan, Männedorf, Switzerland) that can accommodate 384 discreet fermentations and sample automatically at programmable intervals. Fermentation vessels had custom-made airlocks that allowed sampling through a silicon septum and were mixed at inoculation and briefly before sampling with magnetic stir bars. Viognier-Marsanne juice was pressed from solids to alleviate the complexity of changing the juice:solid ratio by dilution in fermentations typically undertaken with most red wines. Temperature was regulated to 17 *°*C by a water bath that housed the fermentation flasks. 25 mL of juice (ameliorated with dilution or addition of FERMAID^®^ O) was inoculated with 5 x 10^6^ yeast cells mL^−1^ (either EC1118™ or Lalvin R2™). After 24 hours, experimental fermentations treated with lactic acid bacteria (LAB+) were inoculated with 0.25 mL of cultured VP41. Control (LAB-) fermentations had 0.25 mL sterile Viognier-Marsanne juice added. Fermentations were performed in triplicate. This combination of treatments of yeast (Lalvin EC1118™, Lalvin R2™ or none), dilution (16, 14.5, 13.5 or 12.5 Bé), nutrient addition (0, 200 or 400 mg L^−1^) and LAB (+/-) resulted in 216 unique fermentations. During fermentation, sugars (glucose, fructose) and malic acid consumption were monitored using modified commercial enzymatic kits (Megazyme, Bray, Ireland) as described in (Walker et al., 2014) and (Jiang, Sumby, Sundstrom, Grbin, & Jiranek, 2018).

### 2.3 Determination of wine composition by HPLC and GC-MS

Major yeast metabolites, organic acids (malic, succinic, acetic), glucose, fructose, glycerol and ethanol were determined from terminal fermentation samples by HPLC according to (Lin, Boss, Walker, Sumby, Grbin, & Jiranek, 2020).

Terminal fermentation samples were also analysed by solid phase micro-extraction (SPME)-GC/MS according to Hranilovic et al. (2018) with extraction and chromatographic condition as outlined by Boss and coworkers (2015). Samples of un-inoculated juices, subjected to the same experimental conditions were also analysed to decipher between juice and fermentation related volatiles. The concentrations of volatiles measured in unfermented juices were very low or below detection (data not shown).

### 2.4 Statistical analysis

Statistical analysis was undertaken with GraphPad Prism v 7.02 (GraphPad Software, La Jolla, CA, USA) and XLSTAT (Addinsoft, New York, NY, USA). Data is reported as the mean values with standard deviation and one way analysis of variance (ANOVA) with Fisher’s least significant difference (LSD) multiple comparison test (p < 0.05) to determine statistical significance. Principal component analysis (PCA) was conducted using The Unscrambler X v10.1 (CAMO Software, Oslo, Norway) and utilised standard scores derived from the volatile compound concentrations measured in each wine sample.

## 3. Results and Discussion

### 3.1 Dilution and addition of nutrients reduced alcoholic and malolactic fermentation duration

Both alcoholic (AF) and malolactic fermentation (MLF) duration were shortened in diluted juices, as is expected where initial sugar concentrations were reduced (AF, Fig. 1 and 2; MF, data not shown). For instance, AF of juice diluted from 16 Bé to 12.5 Bé was reduced in duration with the use of Lalvin EC1118™ by 126 hours, or 42% (298 vs 172 h). AF of undiluted juices by Lalvin R2™ was slightly longer (in comparison to Lalvin EC1118™), however, dilution resulted in a similar reduction of fermentation duration of 121 hours or 34% (355 v 234 h). Dilution to 13.5 Bé or below also allowed the completion of MLF within 45 days in wines fermented by either yeast, whereas MLF of higher Baumé juices failed to complete in this time frame, leaving residual malic acid (>1.0 g L^−1^) (Supp. Tab. 1). The very slow MLF in these conditions is thought to be particular to this juice, perhaps due to limitation of a micronutrient or presence of an inhibitor.

**Figure 1.**
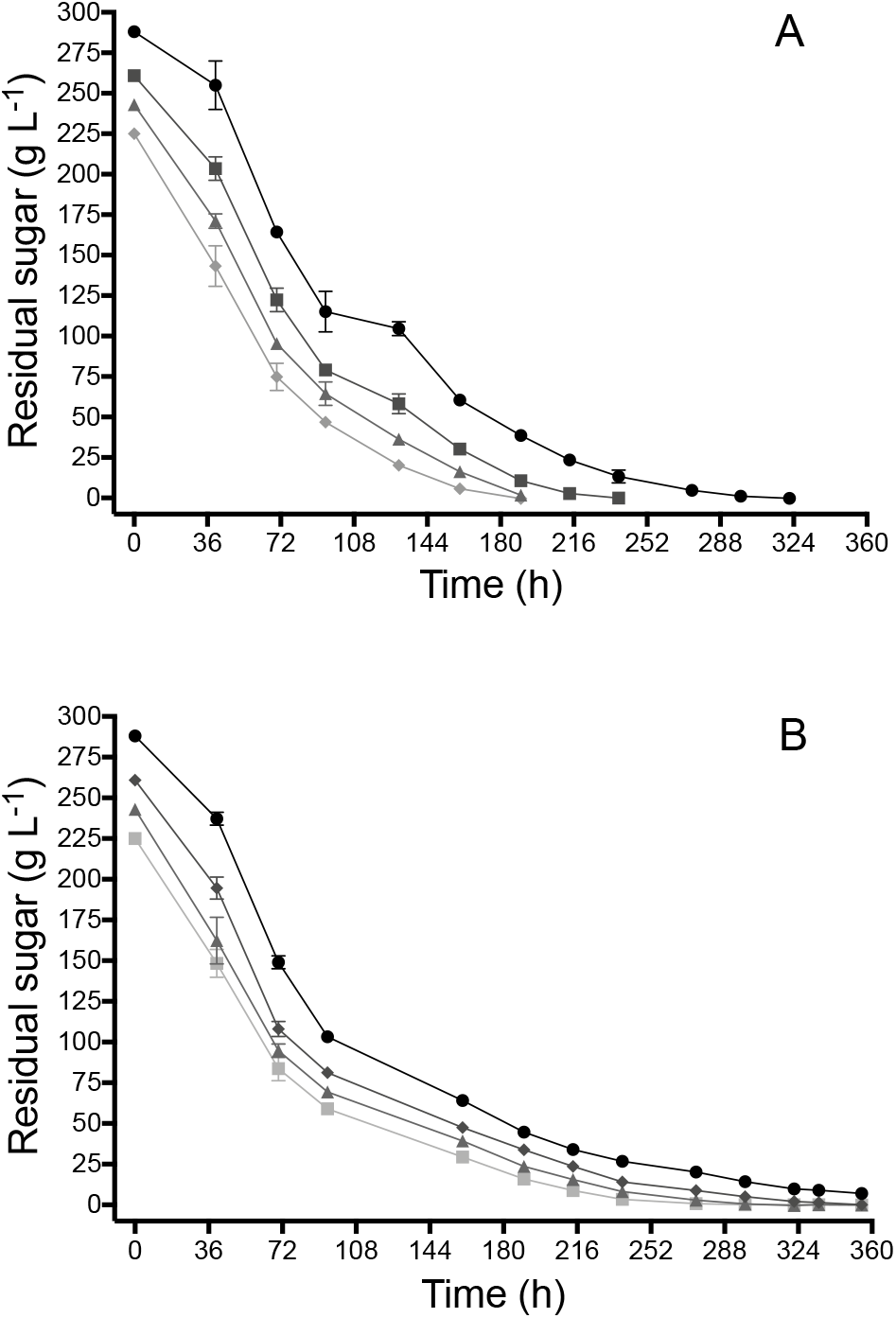
Sugar catabolism by Lalvin EC1118™ (A) or Lalvin R2™ (B) of 16 Bé juice (●) and juices diluted with water to 14.5 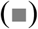, 13.5 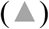 and 12.5 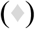 Bé without the addition of malolactic bacteria or complex organic nutrient. Data presented are the average of triplicate fermentations, with the error bars representing the standard deviation.

**Figure 2.**
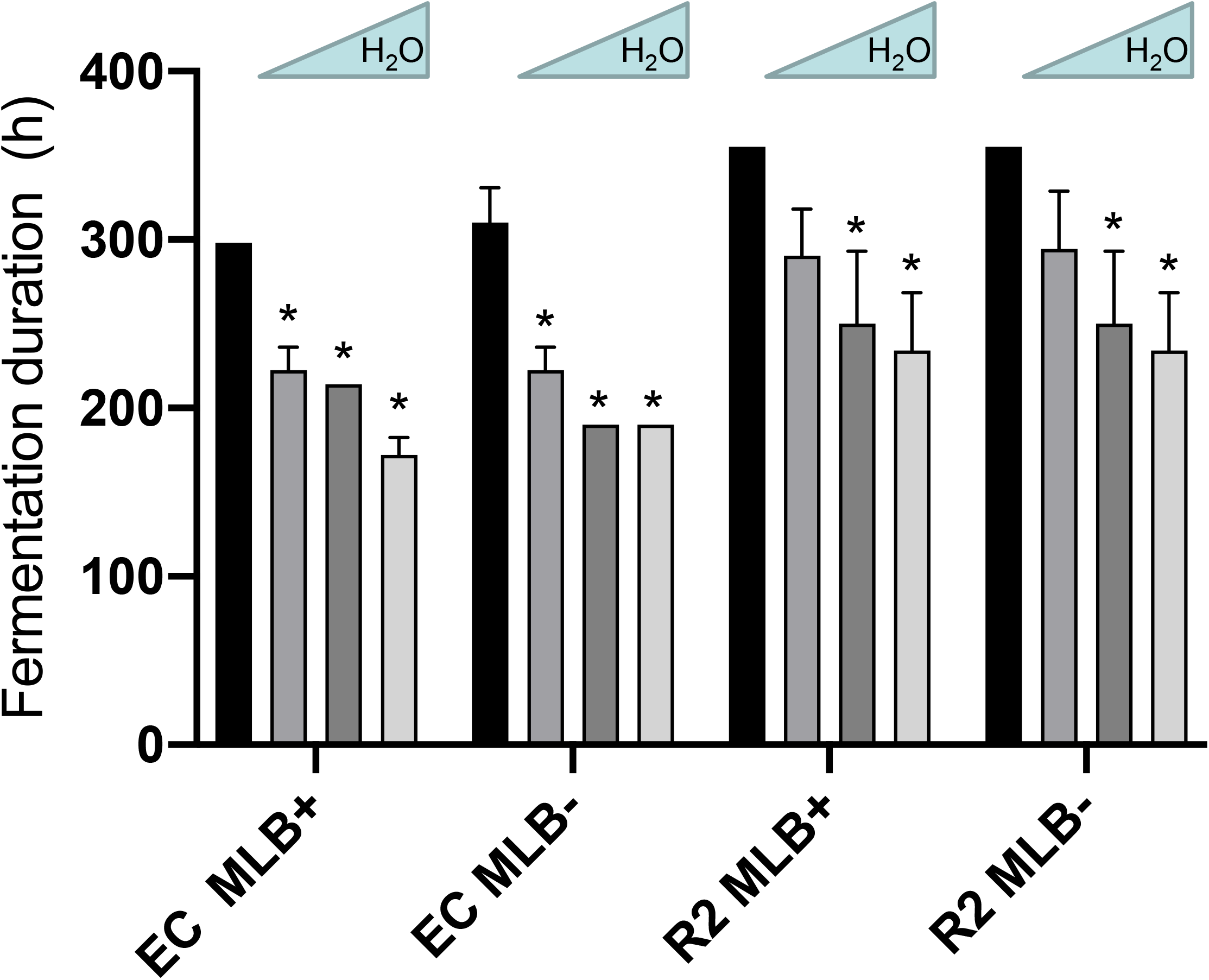
Alcoholic fermentation duration of Lalvin EC1118™ (EC) and Lalvin R2™ (R2), with (MLB+) and without (MLB-) malolactic bacteria. 16 Bé juice was either fermented neat (■), or diluted with water to 14.5 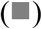, 13.5 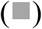 or 12.5 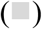 Bé. Data presented is the average of triplicate fermentations and includes standard deviations. *Significantly different to fermentation duration of 16 Bé juice within the set (same yeast and MLB treatment), one way ANOVA, p < 0.05.

The effect on fermentation dynamics of diluted juices with nutrient addition was also evaluated, this was in an effort to alleviate effects, if any, of micronutrient dilution. Additions were made at an industry standard rate, as recommended by the manufacturer (200 mg L^−1^) and also at a higher rate (400 mg L^−1^). The addition of nutrient had no significant impact on AF duration by Lalvin EC1118™ in any juice (data not shown), however, fermentations by Lalvin R2 ™, were up to 76 hours (26%) shorter (Fig. 3). No difference in fermentation dynamics between juices supplemented at either nutrient rate was observed, except in 13.5 Bé juice. Moreover, when either 200 or 400 mg L^−1^ of nutrient was added to 16 Bé juice, fermentation was reduced by 5 hours or 48 hours in 14.5 or 12.5 Bé juices, respectively (data not shown). Whereas addition of 400 mg L^−1^ of nutrient added to 13.5 Bé juices reduced fermentation by 76 hours in comparison to 52 hours when only 200mg L^−1^ of nutrient was added and malolactic acid bacteria (MLB) were present (Fig. 3). The presence of MLB also had no effect on AF duration (Fig. 3).

**Figure 3.**
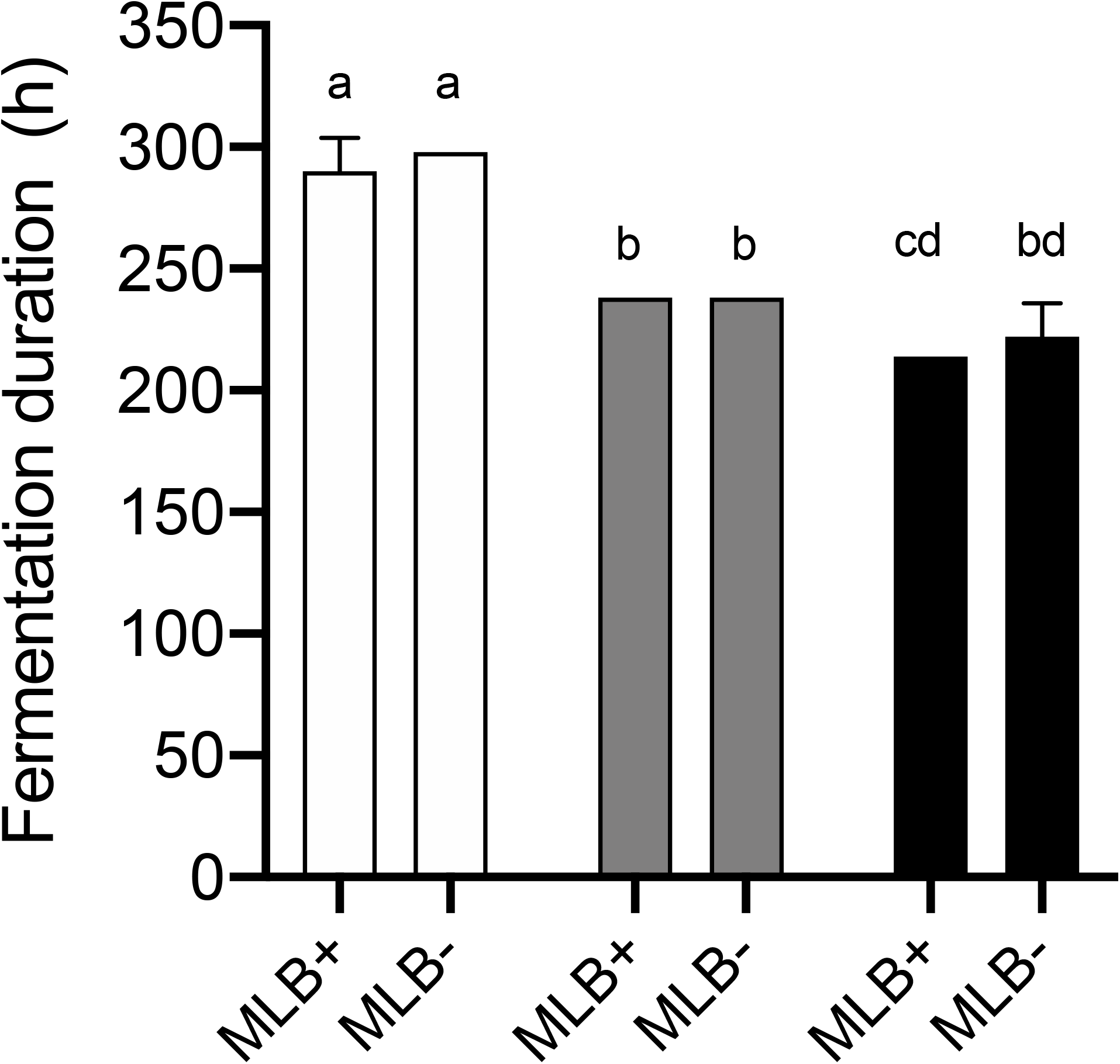
Alcoholic fermentation duration of Lalvin R2™, in juice diluted to 13.5 Bé with (MLB+) and without (MLB-) malolactic bacteria and with the addition of complex organic nutrient at 0 (□), 200 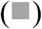 or 400 (■) mg L^−1^. Data presented is the average of triplicate fermentations and includes standard deviations. Values with different letters are significantly different, one way ANOVA, p < 0.05.

The addition of nutrient shortened MLF. Juice diluted to 13.5 Bé with 200 or 400 mg L^−1^ of nutrient and with AF undertaken by Lalvin EC1118™ completed MLF in 47 hours less (4.7% reduction) than when no nutrient was added (Supplementary Table 1).

### 3.2 Dilution and nutrient addition changed the chemical composition of wines

The effect of juice dilution on volatile compounds and major yeast metabolites of the resulting wines was examined. Similar to other studies (Schelezki et al., 2020; Schelezki, Suklje, et al., 2018), the dilution of juices modified the final concentrations of many compounds. Of the 38 volatile compounds analysed, 22 were significantly different to the control in at least three treatments (Table 1) with 19 influenced by dilution and 17 by nutrient addition. In almost all cases, dilution of juices resulted in a reduction in the concentration of volatiles, presumably by dilution of juice-derived precursors that arise from major metabolic pathways such as glycolysis. This is outcome is similar to that seen by Schelezki et al. (2020) where many compounds were reported to decrease upon addition of water to Shiraz juice prior to fermentation.

**Table 1.**
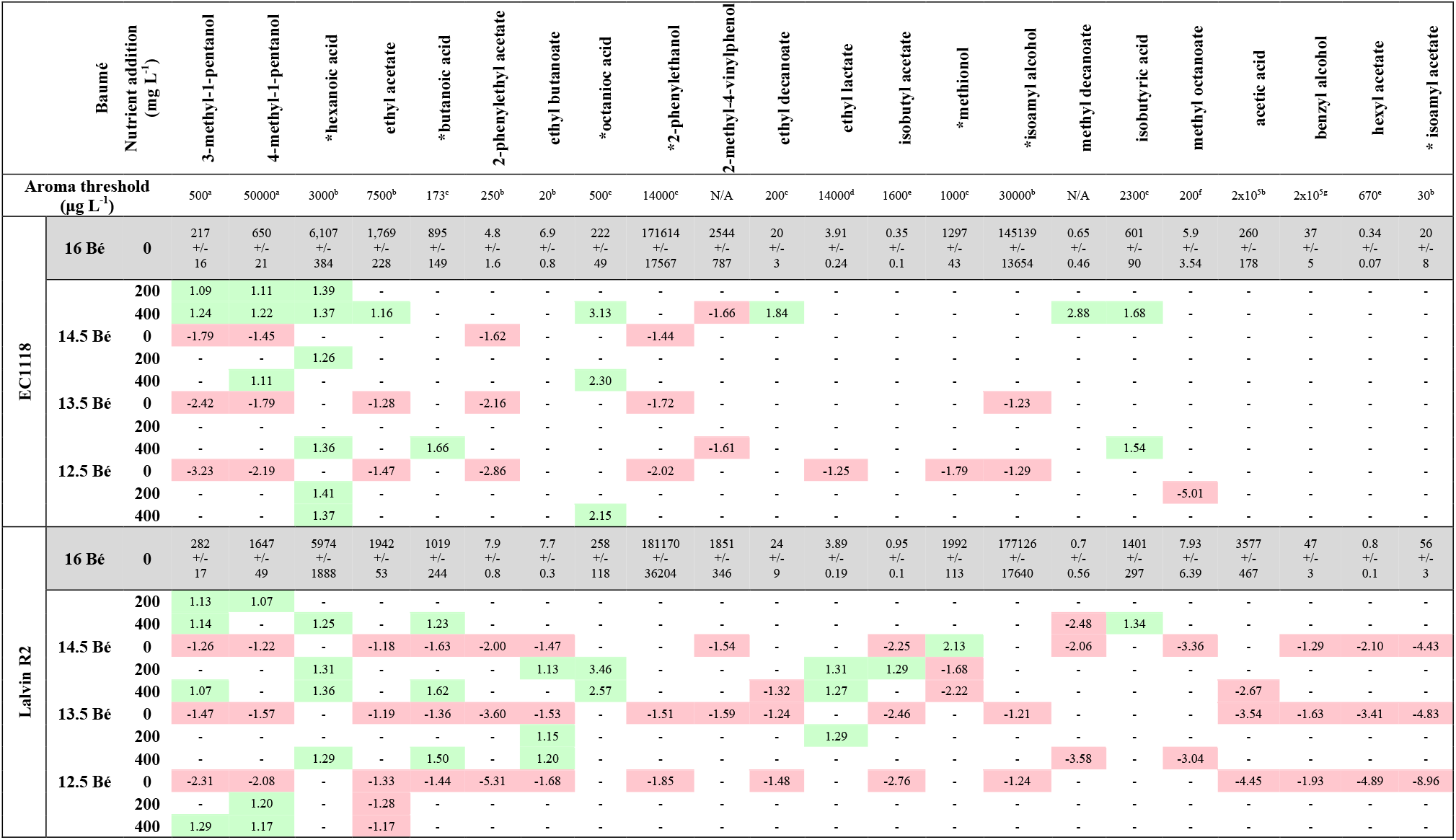
Volatiles detected in final wines that were significantly different (one way ANOVA, p < 0.05) to the matched control (no nutrient added 16 Bé juice fermented by either Lalvin EC1118™ or Lalvin R2 ™ with no exogenous nutrient added, or where nutrients were added comparison was to the same Be juice with no nutrients added). Where a significant difference was detected, the fold change difference of treatment to control is shown. Volatiles that increased upon dilution are shaded in green and decreased in red. – no significant difference. Actual values (μg L^−1^) are shown for undiluted juices. Aroma threshold as reported in ^a^Moreno et al. (2005), ^b^Guth (1997), ^c^Ferreira et al. (2000), ^d^Peinado, Moreno, Bueno, Moreno, & Mauricio (2004); Salo (1970), ^e^Etievant (1991), ^f^Takeoka et al. (1989), ^g^Gomez-Miguez, Cacho, Ferreira, Vicario, & Heredia (2007). *Volatiles detected above the aroma threshold in at least one treatment.

2-Phenylethanol, described as contributing aromas of rose, honey and spice, was detected in all fermentations above its aroma threshold, 14 mg L^−1^ (Ferreira, Lopez, & Cacho, 2000), and with almost all dilutions was significantly reduced (70 - 49% or −1.44 to −2.02 Fold change (FC)). Its ester, 2-phenylethyl acetate was also reduced with dilution (62 – 19% or −1.62 to −5.31 FC). In undiluted fermentations, isoamyl alcohol and, in fermentations conducted with Lalvin R2™, its ester, isoamyl acetate were also detected above their aroma thresholds. Isoamyl acetate could be reduced to below its aroma threshold with any of the trialled dilutions (Fig. 4). Typically, isoamyl acetate was reduced to around 20% of the control (6 +/-3 to 13 +/- 9 vs 56 +/- 3 μg L^−1^, Supp. Tab. 2). This ester can contribute an overpowering banana aroma, especially in white wines, and as such, a dilution strategy (where grapes have an elevated Bé) may represent an option for desirable flavour modification during winemaking.

**Figure 4.**
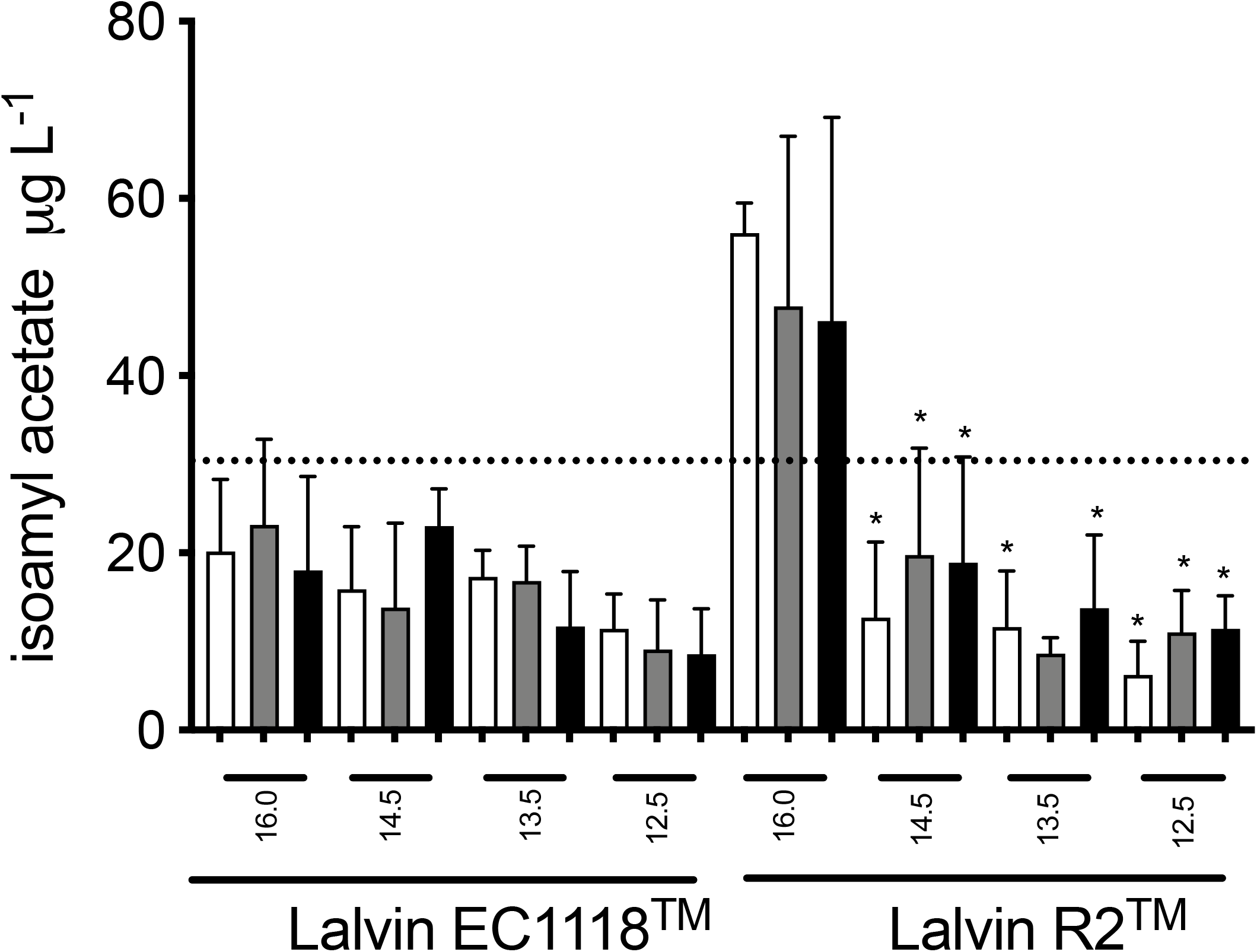
Isoamyl acetate measured in wines (GC-MS) fermented by Lalvin EC1118™ or Lalvin R2 ™ with no malolactic bacteria added. Wines were made from juice with initial Bé of 16 or diluted with water to 14.5, 13.5 or 12.5. Nutrient was also added to juices; 0 (□), 200 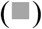 or 400 (■) mg L^−1^. Dashed line represents aroma threshold of isoamyl acetate (30 μg L^−1^ (Guth, 1997)). *significantly different to 16.0 Bé with no nutrient added (for the same yeast), one way ANOVA, p < 0.05).

Acetic acid was also reduced in diluted fermentations conducted with Lalvin R2™, for instance, 3.57 mg L^−1^ was measured in wines using undiluted juice, whilst dilution to 13.5 Bé dramatically reduced acetic acid concentrations to 0.8 mg L^−1^. Even though the aroma threshold of acetic acid is 200 mg L^−1^ (Ferreira et al., 2000), given that this compound is one of the most common faults in wine, even at low concentrations this small difference could impact wine quality. The primary alcohols 3 and 4-methyl-1-pentanol were significantly reduced in almost all diluted juices. For instance, with the use of Lalvin EC1118™, 3-methyl-1-pentanol was reduced to 31% (−3.23 FC) of the control at the most extreme dilution (12.5 Bé). Aromatically, 3-methyl-1-pentanol is described as contributing earthy and green notes, whereas for 4-methyl-1-pentanol its main descriptor is ‘nutty’, however their aroma detection thresholds are near 500 and 50,000 μg L^−1^, respectively (Moreno, Zea, Moyano, & Medina, 2005), many magnitudes above that detected here. If considering these compounds individually we would expect the contribution here to wine aroma to be minimal. It is however widely accepted that many subtle changes in volatile components could result in detectable differences in overall wine aroma, and the individual effects of compounds on the sensory attributes of wine can be complicated by many factors such as the interaction with other wine compounds (Escudero, Campo, Fariña, Cacho, & Ferreira, 2007). Ethyl acetate was also commonly reduced in wines made from diluted juices with either yeast, whereas many more compounds were detected as decreased in diluted juices fermented with Lalvin R2™ i.e., butanoic acid, ethyl butanoate, 2-methyl-4-vinylphenol, ethyl decanoate, isobutyl acetate, benzyl alcohol and hexyl acetate. The single volatile that increased with dilution of juice in this analysis was methionol (sweet potato aroma), but was only significant for the dilution to 14.5 Bé and with the use of Lalvin R2™.

Some major yeast metabolites were also affected by juice dilution (Supp. Tab. 2 – 4). As expected these decreased in concentration in proportion to juice dilution, for example ratios of malic and succinic acid, glycerol and ethanol decreased (ranging from 0.92 to 0.67, Supp. Tab. 4) to very similar ratios to that of juice dilution (14.5–12.5 Bé being 0.89–0.76; juice:water). This confirms the expectation that the major determinate of the final concentrations of these compounds is the initial concentration of sugars in juice. Only the concentration of acetaldehyde increased when juices were diluted to 13.5 or 14.5 Bé and fermented by Lalvin R2™ (167 and 197% respectively). Pyruvate decarboxylase forms acetaldehyde from pyruvate in the latter stages of glycolysis and then alcohol dehydrogenase reduces it to ethanol (Pronk, Yde Steenema, & Van Dijken, 1996). This reaction importantly regenerates NAD^+^ from NADH. The accumulation of acetaldehyde is influenced by the expression and subsequent activity of alcohol dehydrogenases and particularly by the availability of its cofactor, NADH (Xu, Bao, et al., 2019). Transient accumulation of acetaldehyde has also been linked to decreased activity of NADP-dependent acetaldehyde dehydrogenase, which converts acetaldehyde to acetate (Remize, Andrieu, & Dequin, 2000). This may also explain the reduction in acetic acid accumulation in the present study with fermentation by Lalvin R2™, especially as it is reduced beyond that expected from dilution alone (at 13.5 Bé reduced to 28% and 12.5 Bé to 22% (−3.54 and −4.45 FC) of undiluted juice fermentations, Table 1). Perhaps this indicates that the cumulative effect of juice dilution is a reduction of the activity of alcohol dehydrogenase and/or acetaldehyde dehydrogenase, through modification of the NAD^+^/H or NADP/H pools. Dilution of juices is expected to change the external osmolarity that yeast experience at the beginning of fermentation, and it is well documented that many metabolic processes are affected (Varela & Mager, 1996). These minor adjustments of redox cofactors may reflect how the cell achieves balance under these different initial osmotic conditions and results in changes to compound concentrations, such as acetaldehyde and acetic acid reported here. Interestingly, glycerol, the main compound involved in balancing redox factors in response to osmotic stress, is relatively unaffected, only reducing in proportion to juice dilution (Supp. Tab. 4).

We were also particularly interested to see if nutrient addition could recover volatile concentrations to those similar to undiluted juices, supposedly by re-supplying diluted precursors. Of the volatile compounds that were significantly different with addition of nutrient, the vast majority were increased (44 from 56 data points, Table 1), however these rarely recovered the concentration found in undiluted juices. In some conditions, ethyl lactate could be recovered (Supp. Fig 1). Compounds found to increase with the addition of nutrient in more than one condition included 3 and 4-methyl-1-pentanol (up to 129%), butanoic acid (up to 166%), ethyl butanoate (up to 120%), hexanoic acid (up to 141%) and octanoic acid (up to 346%), ethyl lactate (up to 131%) and isobutyric acid (up to 168%; Table 1). Interestingly the occurrence of fatty acids, hexanoic and octanoic acid was unaffected by juice dilution, but consistently increased with nutrient addition (octanoic: 215–346%, hexanoic: 125–141%). This is akin to that seen by other studies (Torrea, Varela, Ugliano, Ancin-Azpilicueta, Leigh Francis, & Henschke, 2011). Similarly, the fatty acid, butanoic acid, also increased with addition of nutrients, however in contrast to hexanoic and octanoic acids, in wines fermented by Lalvin R2™, dilution did result in reduced butanoic acid. Torrea and colleagues (2003) suggest an increase in medium chain fatty acids could simply be due to a relative increase in fatty acid synthesis due to nitrogen supplementation. Fatty acids are produced during fermentation by the fatty acid synthase (FAS) complex from acetyl-CoA and malonyl-CoA during lipid synthesis (Taylor & Kirsop, 1977) with the main source of acetyl CoA during anaerobiosis being acetic acid (Chen, Siewers, & Nielsen, 2012). A number of factors could influence the activity of the enzymes involved and availability of acetyl-CoA in the cytosol such as the supply of cofactors like NADPH (Bloem, Sanchez, Dequin, & Camarasa, 2016) and availability of nutrients (Chen et al., 2012). The mechanism leading to accumulation of medium chain fatty acids under these conditions is still debated but likely involves premature release from the FAS complex due to feedback inhibition by saturated fatty acids and/or an increase in the fatty acid synthetic pathway (Duffour, Malcorps, & Silcock, 2003; Saerens, Delvaux, Verstrepen, Van Dijck, Thevelein, & Delvaux, 2008). Perhaps saturated fatty acids from the complex nutrient contributed to this release of medium chain fatty acids. These commercial preparations commonly contain inactivated yeast and as such have a component of fatty acids along with amino acids, peptides, proteins, polysaccharides, nucleotides, vitamins (thiamine, biotin, pantothenic acid) and minerals (magnesium and zinc) (Lallemand, 2019; Pozo-Bayón, Andújar-Ortiz, & Moreno-Arribas, 2009). However, yeasts for this purpose are grown with plentiful oxygen, and thus saturated fatty acids should be far exceed the concentration of non-saturated fatty acids thus reducing the possibility of feedback inhibition of the FAS complex. Changes to the redox status of yeast have also been shown to affect volatiles produced by yeast (Bloem et al., 2016). Of particular relevance to this study, Bloem and colleagues (2016) report decreased accumulation of medium chain fatty acids (hexanoic and octanoic) during increased demand for NADPH or NADH. As hexanoic, octanoic and butanoic acids were detected many magnitudes above their aroma threshold (hexanoic: 420 μg L^−1^ (Guth, 1997), octanoic: 500 μg L^−1^ and butanoic: 173 μg L^−1^ (Ferreira et al., 2000)), we expect these difference would translate to a sensorial difference. This group of compounds is commonly described as sweaty and cheesy. The presence of fatty acids have also been shown to translate to increases in their fatty acid ethyl ester, for instance ethyl hexanoate and ethyl butanoate (Saerens et al., 2008; Saerens et al., 2006). In this study, the sensorially desirable esters, ethyl lactate and ethyl butanoate, increased with the addition of nutrient (up to 131 and 120%, respectively in wines fermented by Lalvin R2™). Ethyl lactate is described as fruity and ethyl butanoate as floral, fruity, and strawberry like, but in this study they occurred below their aroma thresholds. Exogenously added amino acids have been suggested to be direct precursors of esters, however studies report variable effects of changes in nutrition upon the ester composition of wine, which may be a reflection of changes in juice composition, yeast strain and amount and timing of additions (Hernández-Orte, Ibarz, Cacho, & Ferreira, 2005; Torrea et al., 2011).

Both 3 and 4-methyl-1-pentanol increase with the addition of nutrient in the order of 9-29%, in comparison to wine made from the juice of the same Bé. The most consistent increases of these compounds occurred with addition of 400 mg L^−1^ nutrient, however the magnitude of increases was always well below that of the effect of the initial dilution, thus not resulting in full recovery of reduced volatiles caused by dilution. An increase in these compounds may simply reflect an increase in flux through glycolysis and/or fatty acid synthesis as the 2-keto acid precursors for their formation might be originally derived from pyruvate, as has been shown possible with synthetic pathways built from a combination of yeast and bacterial genes with the goal of efficient production of these compounds for biofuel (Zhang, Sawaya, Eisenberg, & Liao, 2008). This is supported since the *Saccharomyces cerevisiae* enzyme Adh6p, an alcohol dehydrogenase with a strict specificity for NADPH, is capable of completing the final step in formation of methyl pentanols (Zhang et al., 2008). Other aliphatic alcohols can be formed by yeast via the Ehrlich pathway, however these are limited to 5-carbon chains as determined by amino acid precursors (Hazelwood, Daran, van Maris, Pronk, & Dickinson, 2008). 2-Methyl-4-vinylphenol, methyl octanoate, methyl decanoate, acetic acid and ethyl acetate and methionol were found to decrease with the addition of nutrient in selected conditions (Table 1), with methionol being previously reported to decrease with exogenous additions of nitrogen (ca. 70%; Hernández-Orte et al., 2005).

Few differences in major metabolites were detected with added nutrient (Supp. Tab. 4). The most prominent, as with dilution, was an increase of acetaldehyde. Whether this is simply a reflection of increased flux through the glycolytic pathway or accumulation due to reduced availability of cofactors to drive the activity of alcohol dehydrogenases remains to be proven. Acetaldehyde is generally regarded as undesirable when occurring above 100-125 mg L^−1^, as occurred in this study (120-330 mg L^−1^).

Principal Component Analysis of measured volatiles highlighted the interaction between dilution and yeast strain, with the effect of both of these variables on wine volatile compound composition being explained by both of the first 2 principal components (Fig. 5). Wines made with either yeast and from diluted juices were clearly separated, with wines made with Lalvin EC1118™ located more towards the right and the top of the score plot (i.e. they had more positive PC1 and PC2 values) compared to the comparable juices fermented with Lalvin R2™. The clear separation of these yeasts supports what is widely known in the winemaking industry as Lalvin R2™ can make a significant contribution to wine aroma, thereby it is marketed as “aromatic”, whereas Lalvin EC1118™ is considered “neutral”. Wines also grouped according to their juice dilution with higher Bé cultures being located toward the left and top of the plot compared to those with lower Bé (Fig. 5). No clear trend appears to be associated with nutrient additions, reflecting the differential effects depending upon the compound.

**Figure 5.**
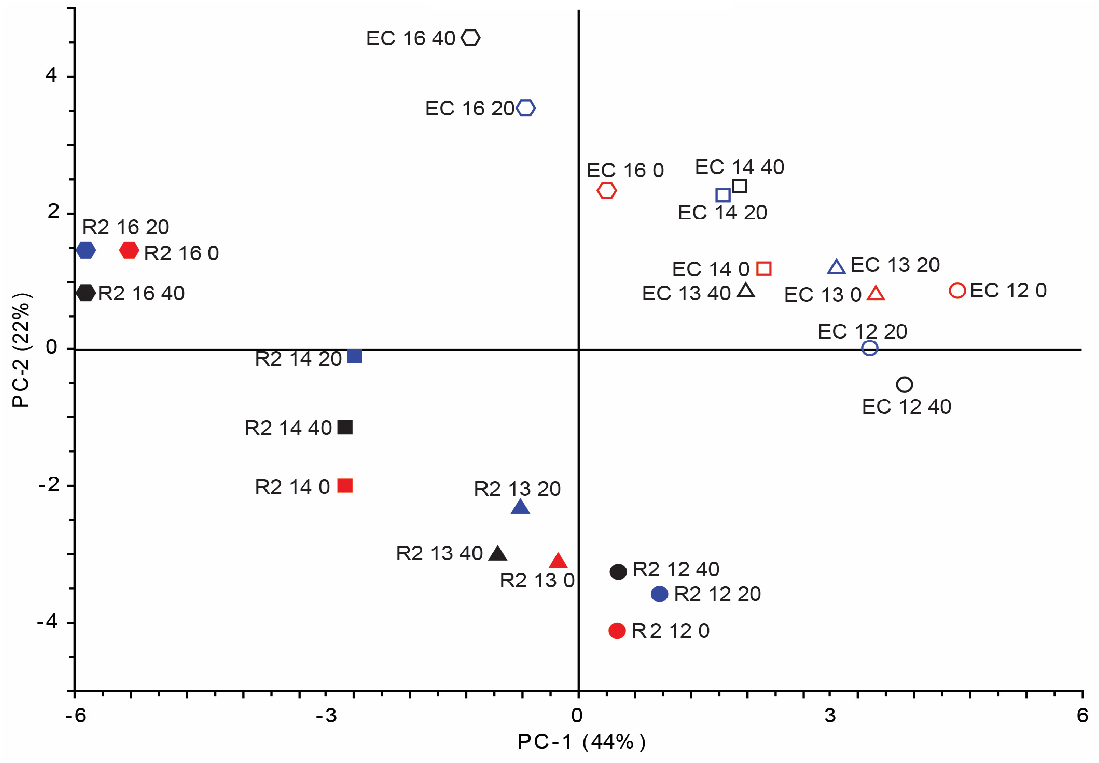
Principal Component Analysis scores plot of volatiles (based on 39 compounds measured by GC-MS) in wines fermented by Lalvin EC1118™ (EC) or Lalvin R2 ™ (R2). Wines made from juice with initial Bé of 16, 14.5 (14), 13.5 (13) and 12.5 (12) with the addition of 0, 200 (20) or 400 (40) mg L^−1^ nutrient. The first two axes are shown (PC-1 and PC-2).

## 4. Conclusions

In this study juice dilution does not impede microbial fermentation, but instead results in reduction of both alcoholic and malolactic fermentation duration and changes to the chemical composition of wines. Nutrient addition was also effective in shortening fermentation duration by Lalvin R2™ by up to 26%. Compounds that were significantly affected in more than 3 treatments by either dilution or nutrient addition and detected above their aroma thresholds were isoamyl acetate, isoamyl alcohol, 2-phenylethanol, methionol and hexanoic, butanoic and octanoic acids. Isoamyl acetate and isoamyl alcohol (banana aroma) and 2-phenylethanol (rose) were reduced by dilution, but the effect upon methionol (sweet potato aroma) varied depending on the treatment. Also the addition of complex organic nutrient, irrespective of dilution rate resulted in increased medium chain fatty acids (hexanoic, butanoic and octanoic characterised by sweaty, cheesy and rancid aromas). The concentrations of 22 volatile compounds were significantly different in three or more treatments. In fact a single treatment resulted in modifications to as many as 15 different volatile compounds and even if these are not above their aroma threshold values, they may act in concert to result in global changes to wine aroma (Escudero et al., 2007). Sensory studies would be of great benefit to determine which of these impacts result in wines that are detectably different to consumers. Depending on the target wine style, these changes may be regarded as a desirable outcome. Winemakers should take into consideration the potential impacts of juice dilution, as well as yeast choice on both processing efficiencies as well as effect on the aroma and flavour of wine.

## Acknowledgements

This work was supported by Wine Australia, Adelaide, South Australia (grant number, UA 1803-2.1). VJ is supported by The Australian Research Council Training Centre for Innovative Wine Production (www.ARCwinecentre.org.au; project number IC170100008), which is funded by the Australian Government with additional support from Wine Australia and industry partners. The University of Adelaide is a member of the Wine Innovation Cluster in Adelaide (http://www.thewaite.org/waite-partners/wine-innovation-cluster/). We also thank Mr Nick van Holst Pellekaan and Dr Jin Zhang for technical assistance during setting up of the fermentations and HPLC analysis and Sue Maffei and Emily Nicholson for assistance with the wine volatile compound analysis. Finally we also thank Lallemand for supply of microbes and complex organic nutrient.

## Conflicts of Interest

We wish to confirm that there are no known conflicts of interest associated with this publication and there has been no significant financial support for this work that could have influenced its outcome.

## Supplementary legends

**Supplementary Table 1.**
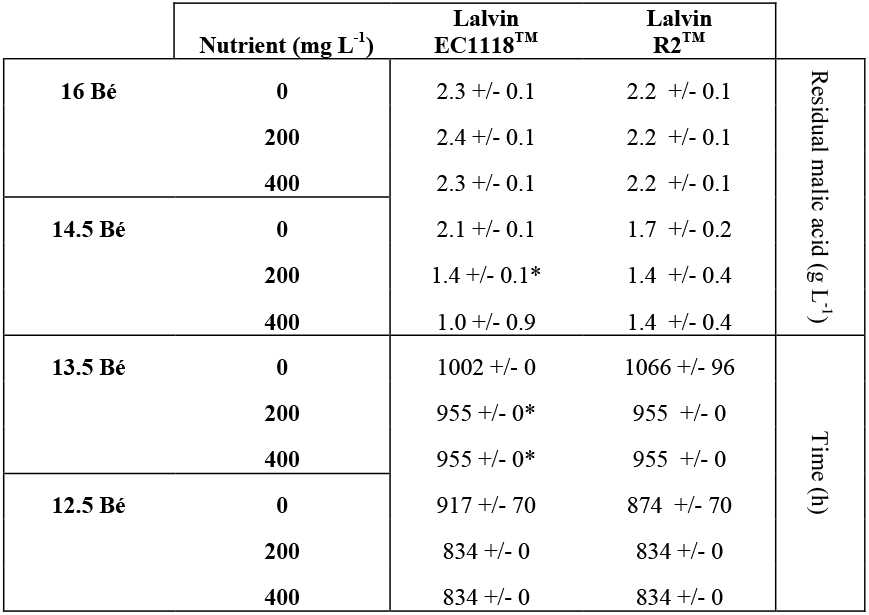
Residual malic acid after 1122 hours or hours taken to metabolise all malic acid (< 0.2 g L^−1^) by VP 41 when co-inoculated with either EC1118™ or Lalvin R2™ in juice at 16 Bé or a range of dilutions with either 0, 200 or 400 mg L^−1^ of complex organic nutrient. Values are the average of triplicates +/- Standard deviations. *Significantly different to no nutrient added (same Bé), Student’s t-test.

**Supplementary Table 2.**
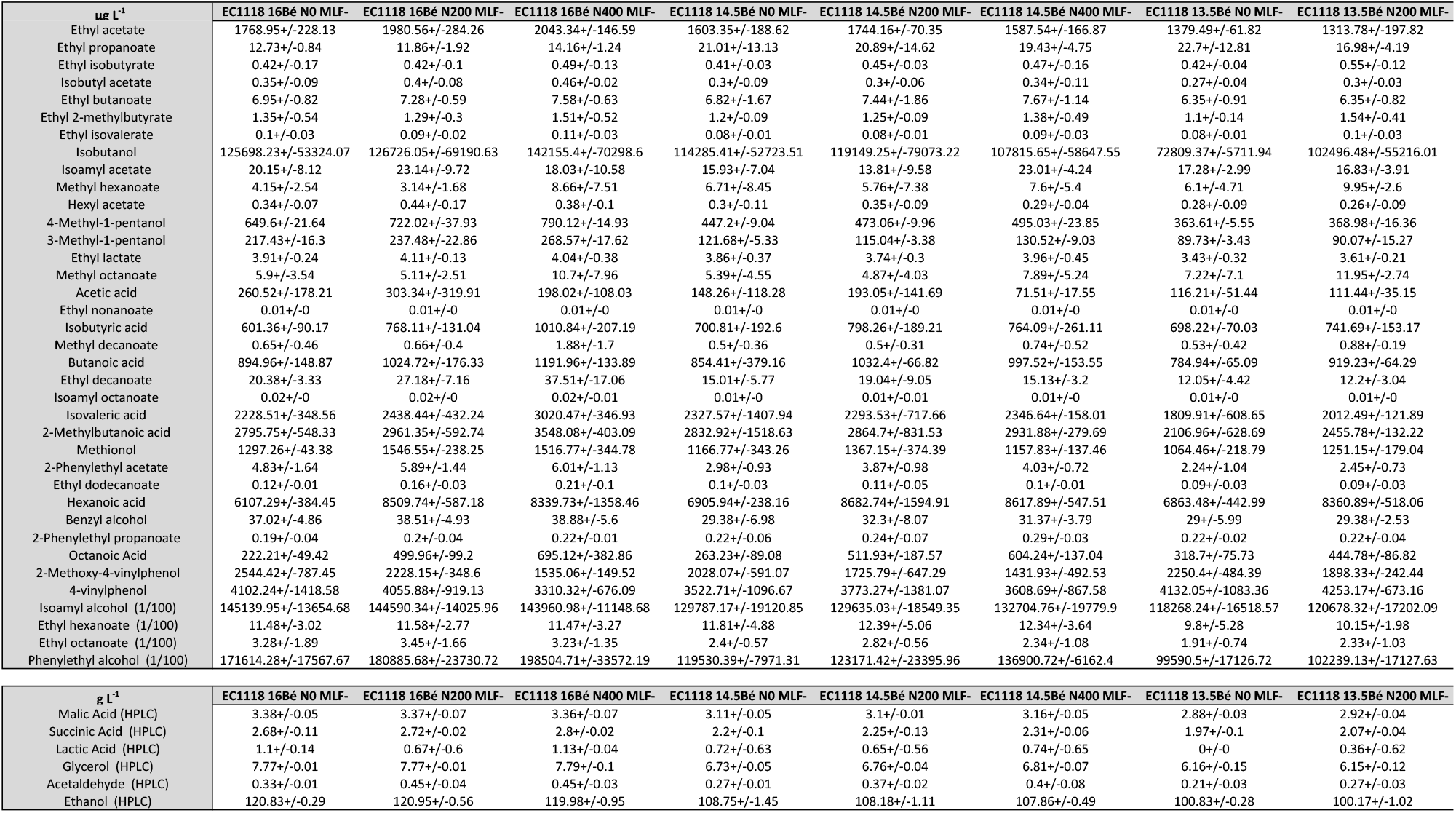
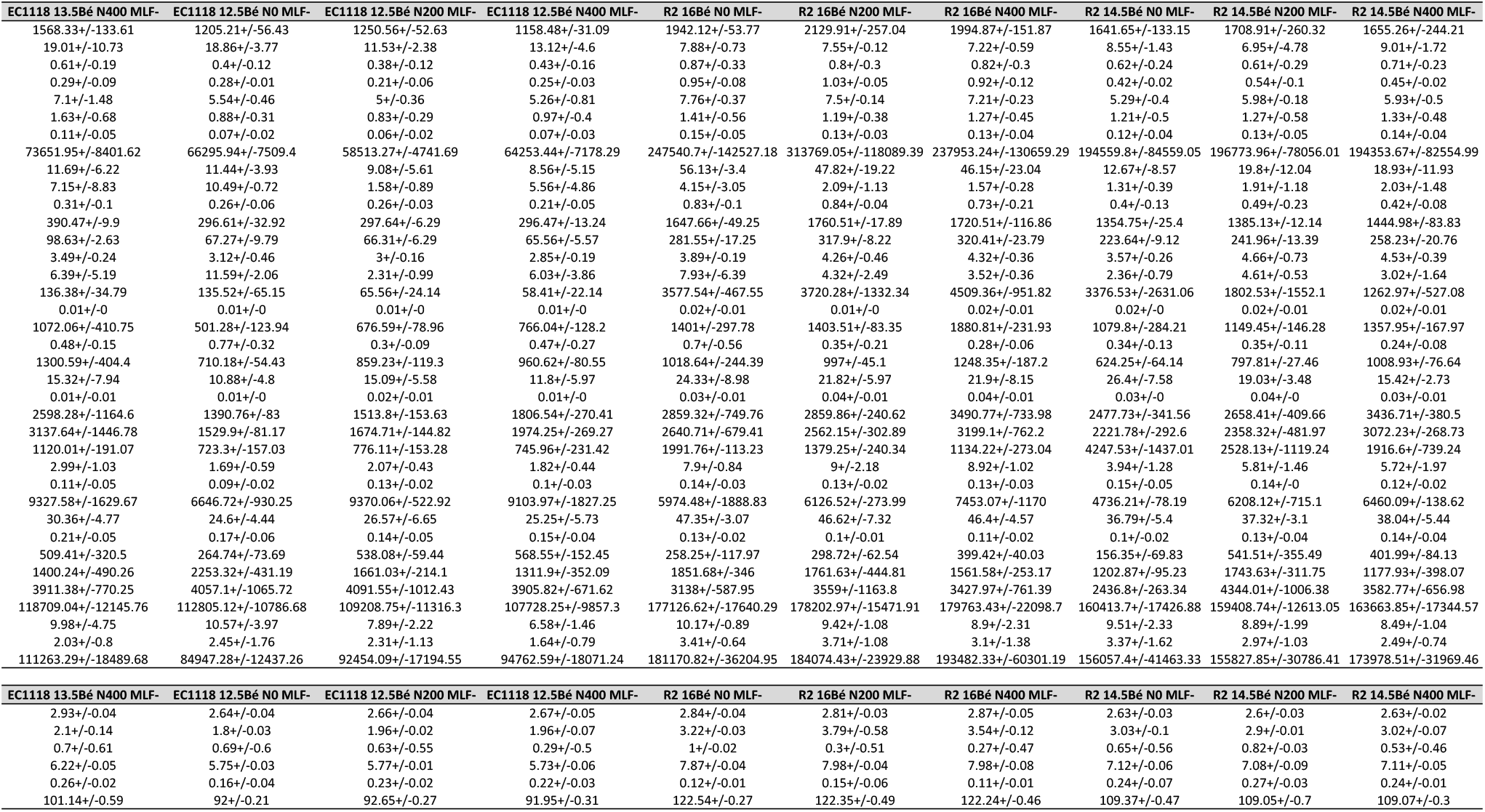
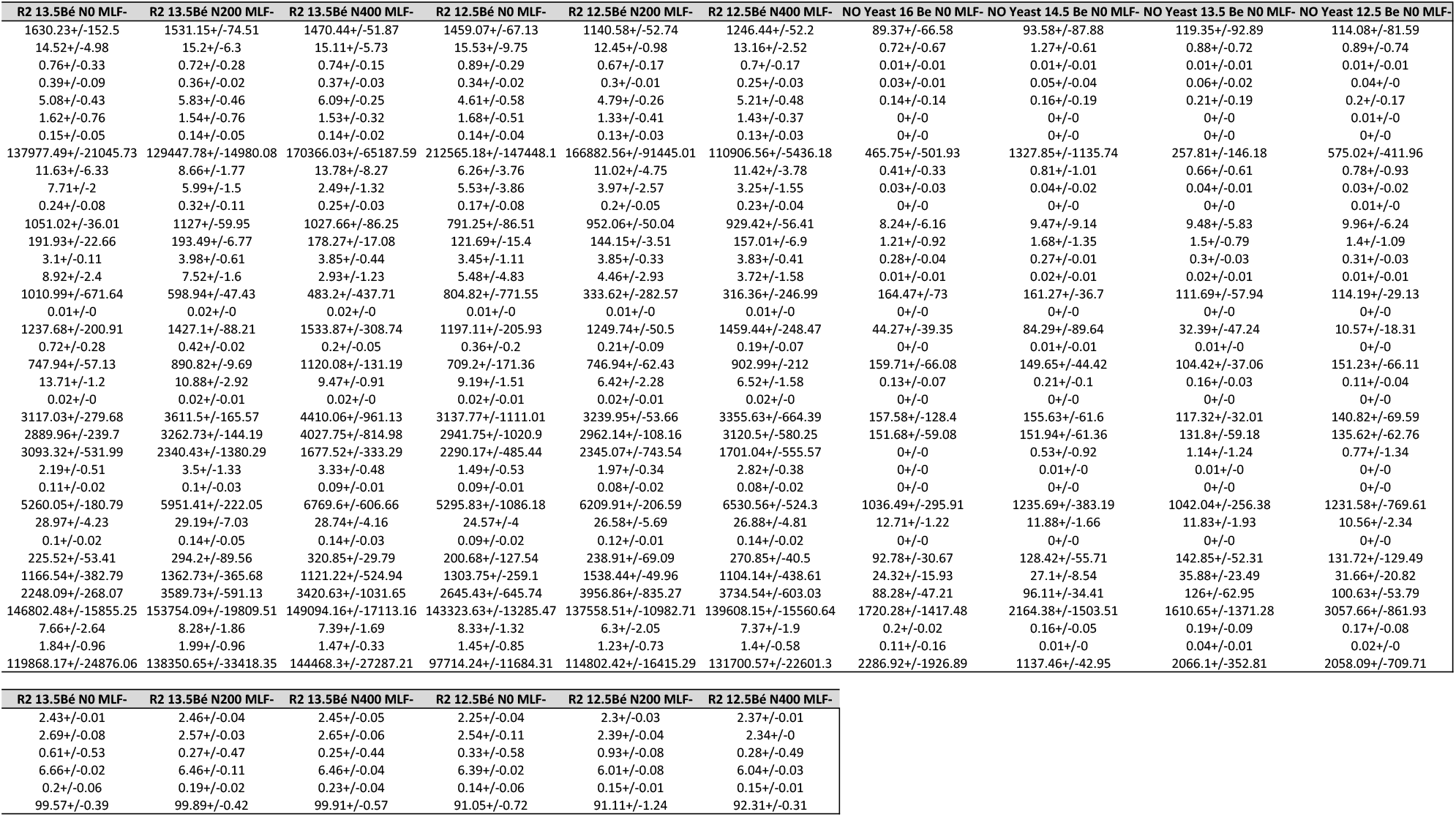
Volatile and major yeast metabolic compounds (μg L^−1^ +/- standard deviations) detected in final wines for all treatments except for those inoculated with lactic acid bacteria for malolactic fermentation. No yeast controls were also analysed, in most cases volatiles were below detection limits, where they were measured they were many magnitudes lower than fermented treatments and thus the data is not shown.

| $\mu\text{g L}^{-1}$ | EC1118 16Bé NO MLF- | EC1118 16Bé N200 MLF- | EC1118 16Bé N400 MLF- | EC1118 14.5Bé NO MLF- | EC1118 14.5Bé N200 MLF- | EC1118 14.5Bé N400 MLF- | EC1118 13.5Bé NO MLF- | EC1118 13.5Bé N200 MLF- |
| --- | --- | --- | --- | --- | --- | --- | --- | --- |
| Ethyl acetate | 1768.95+/-228.13 | 1980.56+/-284.26 | 2043.34+/-146.59 | 1603.35+/-188.62 | 1744.16+/-70.35 | 1587.54+/-166.87 | 1379.49+/-61.82 | 1313.78+/-197.82 |
| Ethyl propanoate | 12.73+/-0.84 | 11.86+/-1.92 | 14.16+/-1.24 | 21.01+/-13.13 | 20.89+/-14.62 | 19.43+/-4.75 | 22.7+/-12.81 | 16.98+/-4.19 |
| Ethyl isobutyrate | 0.42+/-0.17 | 0.42+/-0.1 | 0.49+/-0.13 | 0.41+/-0.03 | 0.45+/-0.03 | 0.47+/-0.16 | 0.42+/-0.04 | 0.55+/-0.12 |
| Isobutyl acetate | 0.35+/-0.09 | 0.4+/-0.08 | 0.46+/-0.02 | 0.3+/-0.09 | 0.3+/-0.06 | 0.34+/-0.11 | 0.27+/-0.04 | 0.3+/-0.03 |
| Ethyl butanoate | 6.95+/-0.82 | 7.28+/-0.59 | 7.58+/-0.63 | 6.82+/-1.67 | 7.44+/-1.86 | 7.67+/-1.14 | 6.35+/-0.91 | 6.35+/-0.82 |
| Ethyl 2-methylbutyrate | 1.35+/-0.54 | 1.29+/-0.3 | 1.51+/-0.52 | 1.2+/-0.09 | 1.25+/-0.09 | 1.38+/-0.49 | 1.1+/-0.14 | 1.54+/-0.41 |
| Ethyl isovalerate | 0.1+/-0.03 | 0.09+/-0.02 | 0.11+/-0.03 | 0.08+/-0.01 | 0.08+/-0.01 | 0.09+/-0.03 | 0.08+/-0.01 | 0.1+/-0.03 |
| Isobutanol | 125698.23+/-53324.07 | 126726.05+/-69190.63 | 142155.4+/-70298.6 | 114285.41+/-52723.51 | 119149.25+/-79073.22 | 107815.65+/-58647.55 | 72809.37+/-5711.94 | 102496.48+/-55216.01 |
| Isoamyl acetate | 20.15+/-8.12 | 23.14+/-9.72 | 18.03+/-10.58 | 15.93+/-7.04 | 13.81+/-9.58 | 23.01+/-4.24 | 17.28+/-2.99 | 16.83+/-3.91 |
| Methyl hexanoate | 4.15+/-2.54 | 3.14+/-1.68 | 8.66+/-7.51 | 6.71+/-8.45 | 5.76+/-7.38 | 7.6+/-5.4 | 6.1+/-4.71 | 9.95+/-2.6 |
| Hexyl acetate | 0.34+/-0.07 | 0.44+/-0.17 | 0.38+/-0.1 | 0.3+/-0.11 | 0.35+/-0.09 | 0.29+/-0.04 | 0.28+/-0.09 | 0.26+/-0.09 |
| 4-Methyl-1-pentanol | 649.6+/-21.64 | 722.02+/-37.93 | 790.12+/-14.93 | 447.2+/-9.04 | 473.06+/-9.96 | 495.03+/-23.85 | 363.61+/-5.55 | 368.98+/-16.36 |
| 3-Methyl-1-pentanol | 217.43+/-16.3 | 237.48+/-22.86 | 268.57+/-17.62 | 121.68+/-5.33 | 115.04+/-3.38 | 130.52+/-9.03 | 89.73+/-3.43 | 90.07+/-15.27 |
| Ethyl lactate | 3.91+/-0.24 | 4.11+/-0.13 | 4.04+/-0.38 | 3.86+/-0.37 | 3.74+/-0.3 | 3.96+/-0.45 | 3.43+/-0.32 | 3.61+/-0.21 |
| Methyl octanoate | 5.9+/-3.54 | 5.11+/-2.51 | 10.7+/-7.96 | 5.39+/-4.55 | 4.87+/-4.03 | 7.89+/-5.24 | 7.22+/-7.1 | 11.95+/-2.74 |
| Acetic acid | 260.52+/-178.21 | 303.34+/-319.91 | 198.02+/-108.03 | 148.26+/-118.28 | 193.05+/-141.69 | 71.51+/-17.55 | 116.21+/-51.44 | 111.44+/-35.15 |
| Ethyl nonanoate | 0.01+/-0 | 0.01+/-0 | 0.01+/-0 | 0.01+/-0 | 0.01+/-0 | 0.01+/-0 | 0.01+/-0 | 0.01+/-0 |
| Isobutyric acid | 601.36+/-90.17 | 768.11+/-131.04 | 1010.84+/-207.19 | 700.81+/-192.6 | 798.26+/-189.21 | 764.09+/-261.11 | 698.22+/-70.03 | 741.69+/-153.17 |
| Methyl decanoate | 0.65+/-0.46 | 0.66+/-0.4 | 1.88+/-1.7 | 0.5+/-0.36 | 0.5+/-0.31 | 0.74+/-0.52 | 0.53+/-0.42 | 0.88+/-0.19 |
| Butanoic acid | 894.96+/-148.87 | 1024.72+/-176.33 | 1191.96+/-133.89 | 854.41+/-379.16 | 1032.4+/-66.82 | 997.52+/-153.55 | 784.94+/-65.09 | 919.23+/-64.29 |
| Ethyl decanoate | 20.38+/-3.33 | 27.18+/-7.16 | 37.51+/-17.06 | 15.01+/-5.77 | 19.04+/-9.05 | 15.13+/-3.2 | 12.05+/-4.42 | 12.2+/-3.04 |
| Isoamyl octanoate | 0.02+/-0 | 0.02+/-0 | 0.02+/-0.01 | 0.01+/-0 | 0.01+/-0.01 | 0.01+/-0 | 0.01+/-0 | 0.01+/-0 |
| Isovaleric acid | 2228.51+/-348.56 | 2438.44+/-432.24 | 3020.47+/-346.93 | 2327.57+/-1407.94 | 2293.53+/-717.66 | 2346.64+/-158.01 | 1809.91+/-608.65 | 2012.49+/-121.89 |
| 2-Methylbutanoic acid | 2795.75+/-548.33 | 2961.35+/-592.74 | 3548.08+/-403.09 | 2832.92+/-1518.63 | 2864.7+/-831.53 | 2931.88+/-279.69 | 2106.96+/-628.69 | 2455.78+/-132.22 |
| Methionol | 1297.26+/-43.38 | 1546.55+/-238.25 | 1516.77+/-344.78 | 1166.77+/-343.26 | 1367.15+/-374.39 | 1157.83+/-137.46 | 1064.46+/-218.79 | 1251.15+/-179.04 |
| 2-Phenylethyl acetate | 4.83+/-1.64 | 5.89+/-1.44 | 6.01+/-1.13 | 2.98+/-0.93 | 3.87+/-0.98 | 4.03+/-0.72 | 2.24+/-1.04 | 2.45+/-0.73 |
| Ethyl dodecanoate | 0.12+/-0.01 | 0.16+/-0.03 | 0.21+/-0.1 | 0.1+/-0.03 | 0.11+/-0.05 | 0.1+/-0.01 | 0.09+/-0.03 | 0.09+/-0.03 |
| Hexanoic acid | 6107.29+/-384.45 | 8509.74+/-587.18 | 8339.73+/-1358.46 | 6905.94+/-238.16 | 8682.74+/-1594.91 | 8617.89+/-547.51 | 6863.48+/-442.99 | 8360.89+/-518.06 |
| Benzyl alcohol | 37.02+/-4.86 | 38.51+/-4.93 | 38.88+/-5.6 | 29.38+/-6.98 | 32.3+/-8.07 | 31.37+/-3.79 | 29+/-5.99 | 29.38+/-2.53 |
| 2-Phenylethyl propanoate | 0.19+/-0.04 | 0.2+/-0.04 | 0.22+/-0.01 | 0.22+/-0.06 | 0.24+/-0.07 | 0.29+/-0.03 | 0.22+/-0.02 | 0.22+/-0.04 |
| Octanoic Acid | 222.21+/-49.42 | 499.96+/-99.2 | 695.12+/-382.86 | 263.23+/-89.08 | 511.93+/-187.57 | 604.24+/-137.04 | 318.7+/-75.73 | 444.78+/-86.82 |
| 2-Methoxy-4-vinylphenol | 2544.42+/-787.45 | 2228.15+/-348.6 | 1535.06+/-149.52 | 2028.07+/-591.07 | 1725.79+/-647.29 | 1431.93+/-492.53 | 2250.4+/-484.39 | 1898.33+/-242.44 |
| 4-vinylphenol | 4102.24+/-1418.58 | 4055.88+/-919.13 | 3310.32+/-676.09 | 3522.71+/-1096.67 | 3773.27+/-1381.07 | 3608.69+/-867.58 | 4132.05+/-1083.36 | 4253.17+/-673.16 |
| Isoamyl alcohol (1/100) | 145139.95+/-13654.68 | 144590.34+/-14025.96 | 143960.98+/-11148.68 | 129787.17+/-19120.85 | 129635.03+/-18549.35 | 132704.76+/-19779.9 | 118268.24+/-16518.57 | 120678.32+/-17202.09 |
| Ethyl hexanoate (1/100) | 11.48+/-3.02 | 11.58+/-2.77 | 11.47+/-3.27 | 11.81+/-4.88 | 12.39+/-5.06 | 12.34+/-3.64 | 9.8+/-5.28 | 10.15+/-1.98 |
| Ethyl octanoate (1/100) | 3.28+/-1.89 | 3.45+/-1.66 | 3.23+/-1.35 | 2.4+/-0.57 | 2.82+/-0.56 | 2.34+/-1.08 | 1.91+/-0.74 | 2.33+/-1.03 |
| Phenylethyl alcohol (1/100) | 171614.28+/-17567.67 | 180885.68+/-23730.72 | 198504.71+/-33572.19 | 119530.39+/-7971.31 | 123171.42+/-23395.96 | 136900.72+/-6162.4 | 99590.5+/-17126.72 | 102239.13+/-17127.63 |

| $\text{g L}^{-1}$ | EC1118 16Bé NO MLF- | EC1118 16Bé N200 MLF- | EC1118 16Bé N400 MLF- | EC1118 14.5Bé NO MLF- | EC1118 14.5Bé N200 MLF- | EC1118 14.5Bé N400 MLF- | EC1118 13.5Bé NO MLF- | EC1118 13.5Bé N200 MLF- |
| --- | --- | --- | --- | --- | --- | --- | --- | --- |
| Malic Acid (HPLC) | 3.38+/-0.05 | 3.37+/-0.07 | 3.36+/-0.07 | 3.11+/-0.05 | 3.1+/-0.01 | 3.16+/-0.05 | 2.88+/-0.03 | 2.92+/-0.04 |
| Succinic Acid (HPLC) | 2.68+/-0.11 | 2.72+/-0.02 | 2.8+/-0.02 | 2.2+/-0.1 | 2.25+/-0.13 | 2.31+/-0.06 | 1.97+/-0.1 | 2.07+/-0.04 |
| Lactic Acid (HPLC) | 1.1+/-0.14 | 0.67+/-0.6 | 1.13+/-0.04 | 0.72+/-0.63 | 0.65+/-0.56 | 0.74+/-0.65 | 0+/-0 | 0.36+/-0.62 |
| Glycerol (HPLC) | 7.77+/-0.01 | 7.77+/-0.01 | 7.79+/-0.1 | 6.73+/-0.05 | 6.76+/-0.04 | 6.81+/-0.07 | 6.16+/-0.15 | 6.15+/-0.12 |
| Acetaldehyde (HPLC) | 0.33+/-0.01 | 0.45+/-0.04 | 0.45+/-0.03 | 0.27+/-0.01 | 0.37+/-0.02 | 0.4+/-0.08 | 0.21+/-0.03 | 0.27+/-0.03 |
| Ethanol (HPLC) | 120.83+/-0.29 | 120.95+/-0.56 | 119.98+/-0.95 | 108.75+/-1.45 | 108.18+/-1.11 | 107.86+/-0.49 | 100.83+/-0.28 | 100.17+/-1.02 |

| EC1118 13.5Bé N400 MLF- | EC1118 12.5Bé NO MLF- | EC1118 12.5Bé N200 MLF- | EC1118 12.5Bé N400 MLF- | R2 16Bé NO MLF- | R2 16Bé N200 MLF- | R2 16Bé N400 MLF- | R2 14.5Bé NO MLF- | R2 14.5Bé N200 MLF- | R2 14.5Bé N400 MLF- |
| --- | --- | --- | --- | --- | --- | --- | --- | --- | --- |
| 1568.33+/-133.61 | 1205.21+/-56.43 | 1250.56+/-52.63 | 1158.48+/-31.09 | 1942.12+/-53.77 | 2129.91+/-257.04 | 1994.87+/-151.87 | 1641.65+/-133.15 | 1708.91+/-260.32 | 1655.26+/-244.21 |
| 19.01+/-10.73 | 18.86+/-3.77 | 11.53+/-2.38 | 13.12+/-4.6 | 7.88+/-0.73 | 7.55+/-0.12 | 7.22+/-0.59 | 8.55+/-1.43 | 6.95+/-4.78 | 9.01+/-1.72 |
| 0.61+/-0.19 | 0.4+/-0.12 | 0.38+/-0.12 | 0.43+/-0.16 | 0.87+/-0.33 | 0.8+/-0.3 | 0.82+/-0.3 | 0.62+/-0.24 | 0.61+/-0.29 | 0.71+/-0.23 |
| 0.29+/-0.09 | 0.28+/-0.01 | 0.21+/-0.06 | 0.25+/-0.03 | 0.95+/-0.08 | 1.03+/-0.05 | 0.92+/-0.12 | 0.42+/-0.02 | 0.54+/-0.1 | 0.45+/-0.02 |
| 7.1+/-1.48 | 5.54+/-0.46 | 5+/-0.36 | 5.26+/-0.81 | 7.76+/-0.37 | 7.5+/-0.14 | 7.21+/-0.23 | 5.29+/-0.4 | 5.98+/-0.18 | 5.93+/-0.5 |
| 1.63+/-0.68 | 0.88+/-0.31 | 0.83+/-0.29 | 0.97+/-0.4 | 1.41+/-0.56 | 1.19+/-0.38 | 1.27+/-0.45 | 1.21+/-0.5 | 1.27+/-0.58 | 1.33+/-0.48 |
| 0.11+/-0.05 | 0.07+/-0.02 | 0.06+/-0.02 | 0.07+/-0.03 | 0.15+/-0.05 | 0.13+/-0.03 | 0.13+/-0.04 | 0.12+/-0.04 | 0.13+/-0.05 | 0.14+/-0.04 |
| 73651.95+/-8401.62 | 66295.94+/-7509.4 | 58513.27+/-4741.69 | 64253.44+/-7178.29 | 247540.7+/-142527.18 | 313769.05+/-118089.39 | 237953.24+/-130659.29 | 194559.8+/-84559.05 | 196773.96+/-78056.01 | 194353.67+/-82554.99 |
| 11.69+/-6.22 | 11.44+/-3.93 | 9.08+/-5.61 | 8.56+/-5.15 | 56.13+/-3.4 | 47.82+/-19.22 | 46.15+/-23.04 | 12.67+/-8.57 | 19.8+/-12.04 | 18.93+/-11.93 |
| 7.15+/-8.83 | 10.49+/-0.72 | 1.58+/-0.89 | 5.56+/-4.86 | 4.15+/-3.05 | 2.09+/-1.13 | 1.57+/-0.28 | 1.31+/-0.39 | 1.91+/-1.18 | 2.03+/-1.48 |
| 0.31+/-0.1 | 0.26+/-0.06 | 0.26+/-0.03 | 0.21+/-0.05 | 0.83+/-0.1 | 0.84+/-0.04 | 0.73+/-0.21 | 0.4+/-0.13 | 0.49+/-0.23 | 0.42+/-0.08 |
| 390.47+/-9.9 | 296.61+/-32.92 | 297.64+/-6.29 | 296.47+/-13.24 | 1647.66+/-49.25 | 1760.51+/-17.89 | 1720.51+/-116.86 | 1354.75+/-25.4 | 1385.13+/-12.14 | 1444.98+/-83.83 |
| 98.63+/-2.63 | 67.27+/-9.79 | 66.31+/-6.29 | 65.56+/-5.57 | 281.55+/-17.25 | 317.9+/-8.22 | 320.41+/-23.79 | 223.64+/-9.12 | 241.96+/-13.39 | 258.23+/-20.76 |
| 3.49+/-0.24 | 3.12+/-0.46 | 3+/-0.16 | 2.85+/-0.19 | 3.89+/-0.19 | 4.26+/-0.46 | 4.32+/-0.36 | 3.57+/-0.26 | 4.66+/-0.73 | 4.53+/-0.39 |
| 6.39+/-5.19 | 11.59+/-2.06 | 2.31+/-0.99 | 6.03+/-3.86 | 7.93+/-6.39 | 4.32+/-2.49 | 3.52+/-0.36 | 2.36+/-0.79 | 4.61+/-0.53 | 3.02+/-1.64 |
| 136.38+/-34.79 | 135.52+/-65.15 | 65.56+/-24.14 | 58.41+/-22.14 | 3577.54+/-467.55 | 3720.28+/-1332.34 | 4509.36+/-951.82 | 3376.53+/-2631.06 | 1802.53+/-1552.1 | 1262.97+/-527.08 |
| 0.01+/-0 | 0.01+/-0 | 0.01+/-0 | 0.01+/-0 | 0.02+/-0.01 | 0.01+/-0 | 0.02+/-0.01 | 0.02+/-0 | 0.02+/-0.01 | 0.02+/-0.01 |
| 1072.06+/-410.75 | 501.28+/-123.94 | 676.59+/-78.96 | 766.04+/-128.2 | 1401+/-297.78 | 1403.51+/-83.35 | 1880.81+/-231.93 | 1079.8+/-284.21 | 1149.45+/-146.28 | 1357.95+/-167.97 |
| 0.48+/-0.15 | 0.77+/-0.32 | 0.3+/-0.09 | 0.47+/-0.27 | 0.7+/-0.56 | 0.35+/-0.21 | 0.28+/-0.06 | 0.34+/-0.13 | 0.35+/-0.11 | 0.24+/-0.08 |
| 1300.59+/-404.4 | 710.18+/-54.43 | 859.23+/-119.3 | 960.62+/-80.55 | 1018.64+/-244.39 | 997+/-45.1 | 1248.35+/-187.2 | 624.25+/-64.14 | 797.81+/-27.46 | 1008.93+/-76.64 |
| 15.32+/-7.94 | 10.88+/-4.8 | 15.09+/-5.58 | 11.8+/-5.97 | 24.33+/-8.98 | 21.82+/-5.97 | 21.9+/-8.15 | 26.4+/-7.58 | 19.03+/-3.48 | 15.42+/-2.73 |
| 0.01+/-0.01 | 0.01+/-0 | 0.02+/-0.01 | 0.01+/-0 | 0.03+/-0.01 | 0.04+/-0.01 | 0.04+/-0.01 | 0.03+/-0 | 0.04+/-0 | 0.03+/-0.01 |
| 2598.28+/-1164.6 | 1390.76+/-83 | 1513.8+/-153.63 | 1806.54+/-270.41 | 2859.32+/-749.76 | 2859.86+/-240.62 | 3490.77+/-733.98 | 2477.73+/-341.56 | 2658.41+/-409.66 | 3436.71+/-380.5 |
| 3137.64+/-1446.78 | 1529.9+/-81.17 | 1674.71+/-144.82 | 1974.25+/-269.27 | 2640.71+/-679.41 | 2562.15+/-302.89 | 3199.1+/-762.2 | 2221.78+/-292.6 | 2358.32+/-481.97 | 3072.23+/-268.73 |
| 1120.01+/-191.07 | 723.3+/-157.03 | 776.11+/-153.28 | 745.96+/-231.42 | 1991.76+/-113.23 | 1379.25+/-240.34 | 1134.22+/-273.04 | 4247.53+/-1437.01 | 2528.13+/-1119.24 | 1916.6+/-739.24 |
| 2.99+/-1.03 | 1.69+/-0.59 | 2.07+/-0.43 | 1.82+/-0.44 | 7.9+/-0.84 | 9+/-2.18 | 8.92+/-1.02 | 3.94+/-1.28 | 5.81+/-1.46 | 5.72+/-1.97 |
| 0.11+/-0.05 | 0.09+/-0.02 | 0.13+/-0.02 | 0.1+/-0.03 | 0.14+/-0.03 | 0.13+/-0.02 | 0.13+/-0.03 | 0.15+/-0.05 | 0.14+/-0 | 0.12+/-0.02 |
| 9327.58+/-1629.67 | 6646.72+/-930.25 | 9370.06+/-522.92 | 9103.97+/-1827.25 | 5974.48+/-1888.83 | 6126.52+/-273.99 | 7453.07+/-1170 | 4736.21+/-78.19 | 6208.12+/-715.1 | 6460.09+/-138.62 |
| 30.36+/-4.77 | 24.6+/-4.44 | 26.57+/-6.65 | 25.25+/-5.73 | 47.35+/-3.07 | 46.62+/-7.32 | 46.4+/-4.57 | 36.79+/-5.4 | 37.32+/-3.1 | 38.04+/-5.44 |
| 0.21+/-0.05 | 0.17+/-0.06 | 0.14+/-0.05 | 0.15+/-0.04 | 0.13+/-0.02 | 0.1+/-0.01 | 0.11+/-0.02 | 0.1+/-0.02 | 0.13+/-0.04 | 0.14+/-0.04 |
| 509.41+/-320.5 | 264.74+/-73.69 | 538.08+/-59.44 | 568.55+/-152.45 | 258.25+/-117.97 | 298.72+/-62.54 | 399.42+/-40.03 | 156.35+/-69.83 | 541.51+/-355.49 | 401.99+/-84.13 |
| 1400.24+/-490.26 | 2253.32+/-431.19 | 1661.03+/-214.1 | 1311.9+/-352.09 | 1851.68+/-346 | 1761.63+/-444.81 | 1561.58+/-253.17 | 1202.87+/-95.23 | 1743.63+/-311.75 | 1177.93+/-398.07 |
| 3911.38+/-770.25 | 4057.1+/-1065.72 | 4091.55+/-1012.43 | 3905.82+/-671.62 | 3138+/-587.95 | 3559+/-1163.8 | 3427.97+/-761.39 | 2436.8+/-263.34 | 4344.01+/-1006.38 | 3582.77+/-656.98 |
| 118709.04+/-12145.76 | 112805.12+/-10786.68 | 109208.75+/-11316.3 | 107728.25+/-9857.3 | 177126.62+/-17640.29 | 178202.97+/-15471.91 | 179763.43+/-22098.7 | 160413.7+/-17426.88 | 159408.74+/-12613.05 | 163663.85+/-17344.57 |
| 9.98+/-4.75 | 10.57+/-3.97 | 7.89+/-2.22 | 6.58+/-1.46 | 10.17+/-0.89 | 9.42+/-1.08 | 8.9+/-2.31 | 9.51+/-2.33 | 8.89+/-1.99 | 8.49+/-1.04 |
| 2.03+/-0.8 | 2.45+/-1.76 | 2.31+/-1.13 | 1.64+/-0.79 | 3.41+/-0.64 | 3.71+/-1.08 | 3.1+/-1.38 | 3.37+/-1.62 | 2.97+/-1.03 | 2.49+/-0.74 |
| 111263.29+/-18489.68 | 84947.28+/-12437.26 | 92454.09+/-17194.55 | 94762.59+/-18071.24 | 181170.82+/-36204.95 | 184074.43+/-23929.88 | 193482.33+/-60301.19 | 156057.4+/-41463.33 | 155827.85+/-30786.41 | 173978.51+/-31969.46 |
| EC1118 13.5Bé N400 MLF- | EC1118 12.5Bé NO MLF- | EC1118 12.5Bé N200 MLF- | EC1118 12.5Bé N400 MLF- | R2 16Bé NO MLF- | R2 16Bé N200 MLF- | R2 16Bé N400 MLF- | R2 14.5Bé NO MLF- | R2 14.5Bé N200 MLF- | R2 14.5Bé N400 MLF- |
| 2.93+/-0.04 | 2.64+/-0.04 | 2.66+/-0.04 | 2.67+/-0.05 | 2.84+/-0.04 | 2.81+/-0.03 | 2.87+/-0.05 | 2.63+/-0.03 | 2.6+/-0.03 | 2.63+/-0.02 |
| 2.1+/-0.14 | 1.8+/-0.03 | 1.96+/-0.02 | 1.96+/-0.07 | 3.22+/-0.03 | 3.79+/-0.58 | 3.54+/-0.12 | 3.03+/-0.1 | 2.9+/-0.01 | 3.02+/-0.07 |
| 0.7+/-0.61 | 0.69+/-0.6 | 0.63+/-0.55 | 0.29+/-0.5 | 1+/-0.02 | 0.3+/-0.51 | 0.27+/-0.47 | 0.65+/-0.56 | 0.82+/-0.03 | 0.53+/-0.46 |
| 6.22+/-0.05 | 5.75+/-0.03 | 5.77+/-0.01 | 5.73+/-0.06 | 7.87+/-0.04 | 7.98+/-0.04 | 7.98+/-0.08 | 7.12+/-0.06 | 7.08+/-0.09 | 7.11+/-0.05 |
| 0.26+/-0.02 | 0.16+/-0.04 | 0.23+/-0.02 | 0.22+/-0.03 | 0.12+/-0.01 | 0.15+/-0.06 | 0.11+/-0.01 | 0.24+/-0.07 | 0.27+/-0.03 | 0.24+/-0.01 |
| 101.14+/-0.59 | 92+/-0.21 | 92.65+/-0.27 | 91.95+/-0.31 | 122.54+/-0.27 | 122.35+/-0.49 | 122.24+/-0.46 | 109.37+/-0.47 | 109.05+/-0.7 | 109.07+/-0.3 |

| R2 13.5Bé N0 MLF- | R2 13.5Bé N200 MLF- | R2 13.5Bé N400 MLF- | R2 12.5Bé N0 MLF- | R2 12.5Bé N200 MLF- | R2 12.5Bé N400 MLF- | NO Yeast 16 Be N0 MLF- | NO Yeast 14.5 Be N0 MLF- | NO Yeast 13.5 Be N0 MLF- | NO Yeast 12.5 Be N0 MLF- |
| --- | --- | --- | --- | --- | --- | --- | --- | --- | --- |
| 1630.23+/-152.5 | 1531.15+/-74.51 | 1470.44+/-51.87 | 1459.07+/-67.13 | 1140.58+/-52.74 | 1246.44+/-52.2 | 89.37+/-66.58 | 93.58+/-87.88 | 119.35+/-92.89 | 114.08+/-81.59 |
| 14.52+/-4.98 | 15.2+/-6.3 | 15.11+/-5.73 | 15.53+/-9.75 | 12.45+/-0.98 | 13.16+/-2.52 | 0.72+/-0.67 | 1.27+/-0.61 | 0.88+/-0.72 | 0.89+/-0.74 |
| 0.76+/-0.33 | 0.72+/-0.28 | 0.74+/-0.15 | 0.89+/-0.29 | 0.67+/-0.17 | 0.7+/-0.17 | 0.01+/-0.01 | 0.01+/-0.01 | 0.01+/-0.01 | 0.01+/-0.01 |
| 0.39+/-0.09 | 0.36+/-0.02 | 0.37+/-0.03 | 0.34+/-0.02 | 0.3+/-0.01 | 0.25+/-0.03 | 0.03+/-0.01 | 0.05+/-0.04 | 0.06+/-0.02 | 0.04+/-0 |
| 5.08+/-0.43 | 5.83+/-0.46 | 6.09+/-0.25 | 4.61+/-0.58 | 4.79+/-0.26 | 5.21+/-0.48 | 0.14+/-0.14 | 0.16+/-0.19 | 0.21+/-0.19 | 0.2+/-0.17 |
| 1.62+/-0.76 | 1.54+/-0.76 | 1.53+/-0.32 | 1.68+/-0.51 | 1.33+/-0.41 | 1.43+/-0.37 | 0+/-0 | 0+/-0 | 0+/-0 | 0.01+/-0 |
| 0.15+/-0.05 | 0.14+/-0.05 | 0.14+/-0.02 | 0.14+/-0.04 | 0.13+/-0.03 | 0.13+/-0.03 | 0+/-0 | 0+/-0 | 0+/-0 | 0+/-0 |
| 137977.49+/-21045.73 | 129447.78+/-14980.08 | 170366.03+/-65187.59 | 212565.18+/-147448.1 | 166882.56+/-91445.01 | 110906.56+/-5436.18 | 465.75+/-501.93 | 1327.85+/-1135.74 | 257.81+/-146.18 | 575.02+/-411.96 |
| 11.63+/-6.33 | 8.66+/-1.77 | 13.78+/-8.27 | 6.26+/-3.76 | 11.02+/-4.75 | 11.42+/-3.78 | 0.41+/-0.33 | 0.81+/-1.01 | 0.66+/-0.61 | 0.78+/-0.93 |
| 7.71+/-2 | 5.99+/-1.5 | 2.49+/-1.32 | 5.53+/-3.86 | 3.97+/-2.57 | 3.25+/-1.55 | 0.03+/-0.03 | 0.04+/-0.02 | 0.04+/-0.01 | 0.03+/-0.02 |
| 0.24+/-0.08 | 0.32+/-0.11 | 0.25+/-0.03 | 0.17+/-0.08 | 0.2+/-0.05 | 0.23+/-0.04 | 0+/-0 | 0+/-0 | 0+/-0 | 0.01+/-0 |
| 1051.02+/-36.01 | 1127+/-59.95 | 1027.66+/-86.25 | 791.25+/-86.51 | 952.06+/-50.04 | 929.42+/-56.41 | 8.24+/-6.16 | 9.47+/-9.14 | 9.48+/-5.83 | 9.96+/-6.24 |
| 191.93+/-22.66 | 193.49+/-6.77 | 178.27+/-17.08 | 121.69+/-15.4 | 144.15+/-3.51 | 157.01+/-6.9 | 1.21+/-0.92 | 1.68+/-1.35 | 1.5+/-0.79 | 1.4+/-1.09 |
| 3.1+/-0.11 | 3.98+/-0.61 | 3.85+/-0.44 | 3.45+/-1.11 | 3.85+/-0.33 | 3.83+/-0.41 | 0.28+/-0.04 | 0.27+/-0.01 | 0.31+/-0.03 | 0.31+/-0.03 |
| 8.92+/-2.4 | 7.52+/-1.6 | 2.93+/-1.23 | 5.48+/-4.83 | 4.46+/-2.93 | 3.72+/-1.58 | 0.01+/-0.01 | 0.02+/-0.01 | 0.02+/-0.01 | 0.01+/-0.01 |
| 1010.99+/-671.64 | 598.94+/-47.43 | 483.2+/-437.71 | 804.82+/-771.55 | 333.62+/-282.57 | 316.36+/-246.99 | 164.47+/-73 | 161.27+/-36.7 | 111.69+/-57.94 | 114.19+/-29.13 |
| 0.01+/-0 | 0.02+/-0 | 0.02+/-0 | 0.01+/-0 | 0.01+/-0 | 0.01+/-0 | 0+/-0 | 0+/-0 | 0+/-0 | 0+/-0 |
| 1237.68+/-200.91 | 1427.1+/-88.21 | 1533.87+/-308.74 | 1197.11+/-205.93 | 1249.74+/-50.5 | 1459.44+/-248.47 | 44.27+/-39.35 | 84.29+/-89.64 | 32.39+/-47.24 | 10.57+/-18.31 |
| 0.72+/-0.28 | 0.42+/-0.02 | 0.2+/-0.05 | 0.36+/-0.2 | 0.21+/-0.09 | 0.19+/-0.07 | 0+/-0 | 0.01+/-0.01 | 0.01+/-0 | 0+/-0 |
| 747.94+/-57.13 | 890.82+/-9.69 | 1120.08+/-131.19 | 709.2+/-171.36 | 746.94+/-62.43 | 902.99+/-212 | 159.71+/-66.08 | 149.65+/-44.42 | 104.42+/-37.06 | 151.23+/-66.11 |
| 13.71+/-1.2 | 10.88+/-2.92 | 9.47+/-0.91 | 9.19+/-1.51 | 6.42+/-2.28 | 6.52+/-1.58 | 0.13+/-0.07 | 0.21+/-0.1 | 0.16+/-0.03 | 0.11+/-0.04 |
| 0.02+/-0 | 0.02+/-0.01 | 0.02+/-0 | 0.02+/-0.01 | 0.02+/-0.01 | 0.02+/-0 | 0+/-0 | 0+/-0 | 0+/-0 | 0+/-0 |
| 3117.03+/-279.68 | 3611.5+/-165.57 | 4410.06+/-961.13 | 3137.77+/-1111.01 | 3239.95+/-53.66 | 3355.63+/-664.39 | 157.58+/-128.4 | 155.63+/-61.6 | 117.32+/-32.01 | 140.82+/-69.59 |
| 2889.96+/-239.7 | 3262.73+/-144.19 | 4027.75+/-814.98 | 2941.75+/-1020.9 | 2962.14+/-108.16 | 3120.5+/-580.25 | 151.68+/-59.08 | 151.94+/-61.36 | 131.8+/-59.18 | 135.62+/-62.76 |
| 3093.32+/-531.99 | 2340.43+/-1380.29 | 1677.52+/-333.29 | 2290.17+/-485.44 | 2345.07+/-743.54 | 1701.04+/-555.57 | 0+/-0 | 0.53+/-0.92 | 1.14+/-1.24 | 0.77+/-1.34 |
| 2.19+/-0.51 | 3.5+/-1.33 | 3.33+/-0.48 | 1.49+/-0.53 | 1.97+/-0.34 | 2.82+/-0.38 | 0+/-0 | 0.01+/-0 | 0.01+/-0 | 0+/-0 |
| 0.11+/-0.02 | 0.1+/-0.03 | 0.09+/-0.01 | 0.09+/-0.01 | 0.08+/-0.02 | 0.08+/-0.02 | 0+/-0 | 0+/-0 | 0+/-0 | 0+/-0 |
| 5260.05+/-180.79 | 5951.41+/-222.05 | 6769.6+/-606.66 | 5295.83+/-1086.18 | 6209.91+/-206.59 | 6530.56+/-524.3 | 1036.49+/-295.91 | 1235.69+/-383.19 | 1042.04+/-256.38 | 1231.58+/-769.61 |
| 28.97+/-4.23 | 29.19+/-7.03 | 28.74+/-4.16 | 24.57+/-4 | 26.58+/-5.69 | 26.88+/-4.81 | 12.71+/-1.22 | 11.88+/-1.66 | 11.83+/-1.93 | 10.56+/-2.34 |
| 0.1+/-0.02 | 0.14+/-0.05 | 0.14+/-0.03 | 0.09+/-0.02 | 0.12+/-0.01 | 0.14+/-0.02 | 0+/-0 | 0+/-0 | 0+/-0 | 0+/-0 |
| 225.52+/-53.41 | 294.2+/-89.56 | 320.85+/-29.79 | 200.68+/-127.54 | 238.91+/-69.09 | 270.85+/-40.5 | 92.78+/-30.67 | 128.42+/-55.71 | 142.85+/-52.31 | 131.72+/-129.49 |
| 1166.54+/-382.79 | 1362.73+/-365.68 | 1121.22+/-524.94 | 1303.75+/-259.1 | 1538.44+/-49.96 | 1104.14+/-438.61 | 24.32+/-15.93 | 27.1+/-8.54 | 35.88+/-23.49 | 31.66+/-20.82 |
| 2248.09+/-268.07 | 3589.73+/-591.13 | 3420.63+/-1031.65 | 2645.43+/-645.74 | 3956.86+/-835.27 | 3734.54+/-603.03 | 88.28+/-47.21 | 96.11+/-34.41 | 126+/-62.95 | 100.63+/-53.79 |
| 146802.48+/-15855.25 | 153754.09+/-19809.51 | 149094.16+/-17113.16 | 143323.63+/-13285.47 | 137558.51+/-10982.71 | 139608.15+/-15560.64 | 1720.28+/-1417.48 | 2164.38+/-1503.51 | 1610.65+/-1371.28 | 3057.66+/-861.93 |
| 7.66+/-2.64 | 8.28+/-1.86 | 7.39+/-1.69 | 8.33+/-1.32 | 6.3+/-2.05 | 7.37+/-1.9 | 0.2+/-0.02 | 0.16+/-0.05 | 0.19+/-0.09 | 0.17+/-0.08 |
| 1.84+/-0.96 | 1.99+/-0.96 | 1.47+/-0.33 | 1.45+/-0.85 | 1.23+/-0.73 | 1.4+/-0.58 | 0.11+/-0.16 | 0.01+/-0 | 0.04+/-0.01 | 0.02+/-0 |
| 119868.17+/-24876.06 | 138350.65+/-33418.35 | 144468.3+/-27287.21 | 97714.24+/-11684.31 | 114802.42+/-16415.29 | 131700.57+/-22601.3 | 2286.92+/-1926.89 | 1137.46+/-42.95 | 2066.1+/-352.81 | 2058.09+/-709.71 |

| R2 13.5Bé N0 MLF- | R2 13.5Bé N200 MLF- | R2 13.5Bé N400 MLF- | R2 12.5Bé N0 MLF- | R2 12.5Bé N200 MLF- | R2 12.5Bé N400 MLF- |
| --- | --- | --- | --- | --- | --- |
| 2.43+/-0.01 | 2.46+/-0.04 | 2.45+/-0.05 | 2.25+/-0.04 | 2.3+/-0.03 | 2.37+/-0.01 |
| 2.69+/-0.08 | 2.57+/-0.03 | 2.65+/-0.06 | 2.54+/-0.11 | 2.39+/-0.04 | 2.34+/-0 |
| 0.61+/-0.53 | 0.27+/-0.47 | 0.25+/-0.44 | 0.33+/-0.58 | 0.93+/-0.08 | 0.28+/-0.49 |
| 6.66+/-0.02 | 6.46+/-0.11 | 6.46+/-0.04 | 6.39+/-0.02 | 6.01+/-0.08 | 6.04+/-0.03 |
| 0.2+/-0.06 | 0.19+/-0.02 | 0.23+/-0.04 | 0.14+/-0.06 | 0.15+/-0.01 | 0.15+/-0.01 |
| 99.57+/-0.39 | 99.89+/-0.42 | 99.91+/-0.57 | 91.05+/-0.72 | 91.11+/-1.24 | 92.31+/-0.31 |

**Supplementary Table 3.**
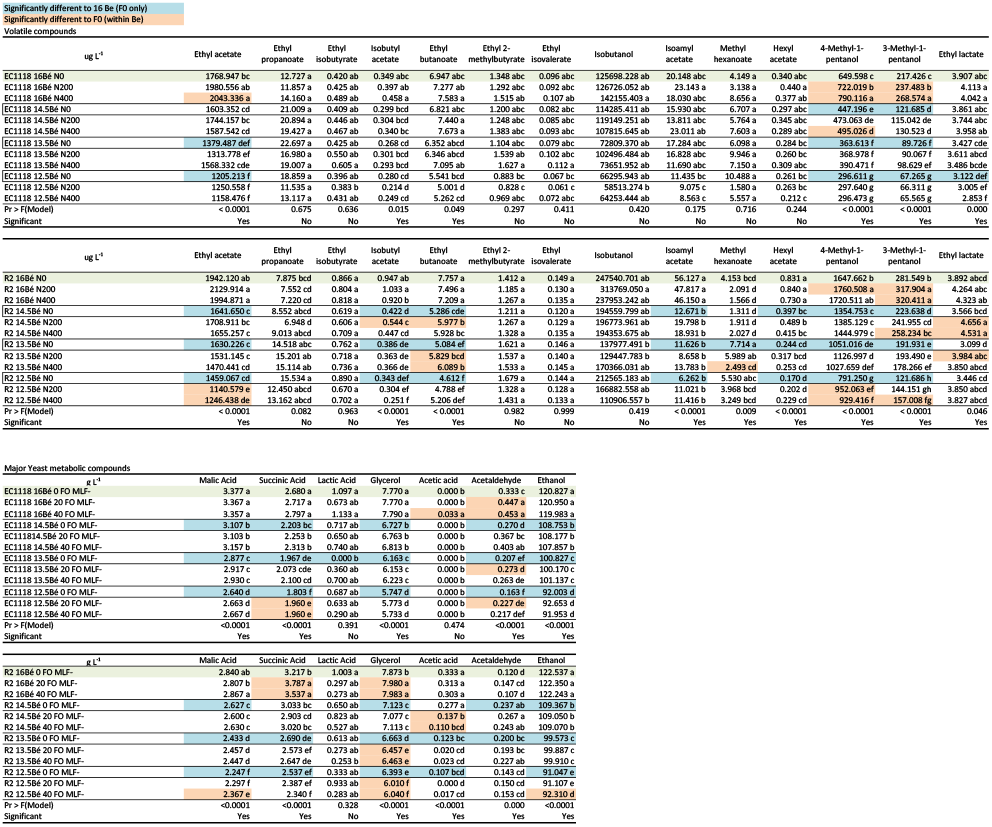
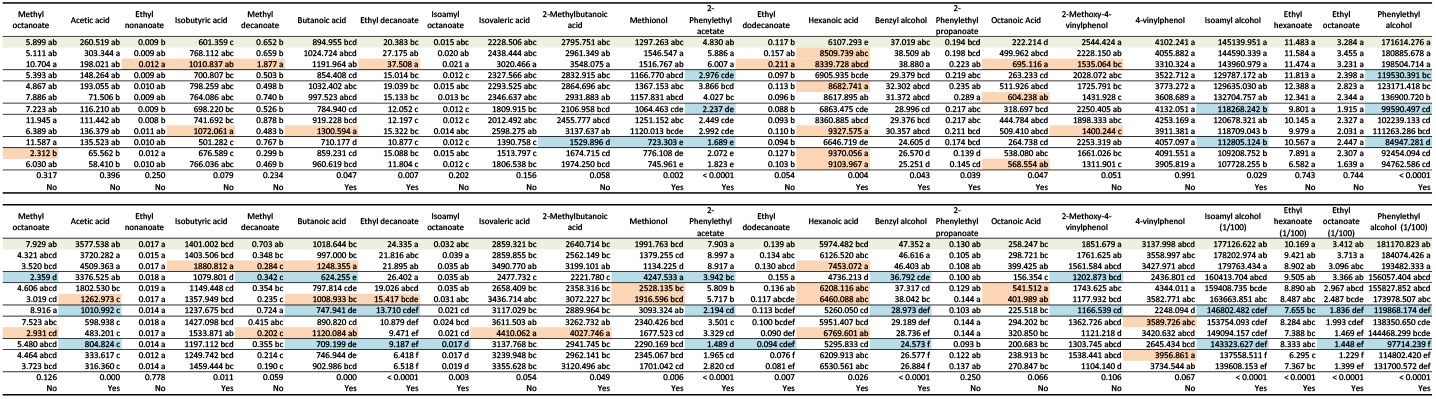
Volatile and major yeast metabolic compounds (volatiles: μg L^−1^ and major metabolites g L^−1^) detected in final wines for all treatments except for those inoculated with lactic acid bacteria for malolactic fermentation. No yeast controls were also analysed, in most cases volatiles were below detection limits, where they were measured they were many magnitudes lower than fermented treatments and thus the data is not shown. Blue highlighted cells indicate those measurements significantly different due to juice dilution (compared to the same yeast at 16 Bé) and orange highlighted cells indicate those significantly different due to nutrient addition (compared to the same yeast and same initial Bé juice). Significant differences were determined by ANOVA and significantly different data are indicated by different letters.

**Supplementary Table 4.**
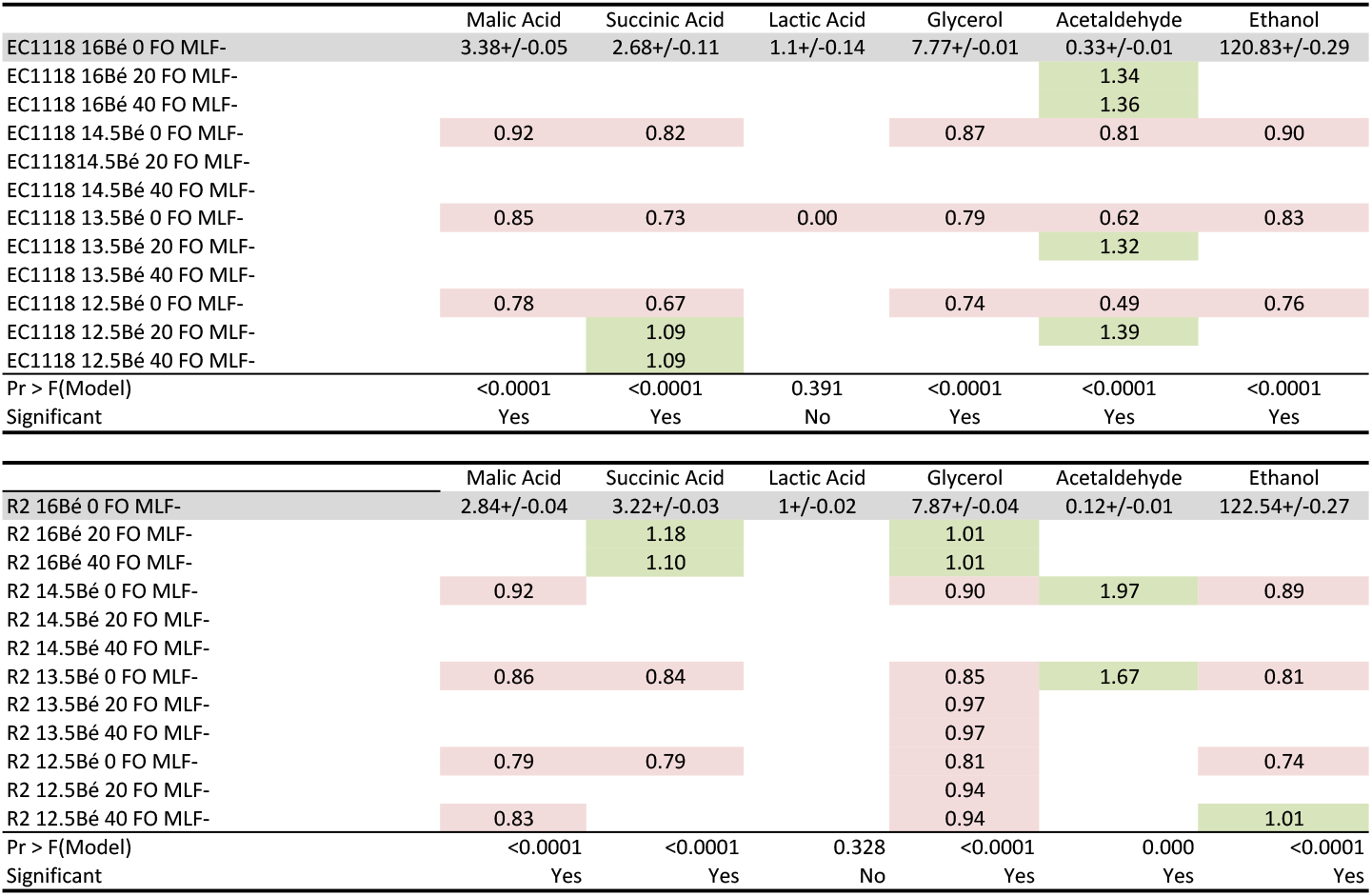
Ratios of significantly different compounds measured by HPLC. Ratios of compounds detected in wines made from diluted juices are in comparison to 16 Bé juices fermented by the same yeast, or in the case of where nutrient is added, to wines made from juices of the same Bé. Increased ratios are highlighted in green and decreased in red. Results for wines made from 16 Bé juices with no nutrient addition are displayed as actual values (g L^−1^) +/- SD. Significant differences were determined by ANOVA.

**Supplementary Figure 1.**
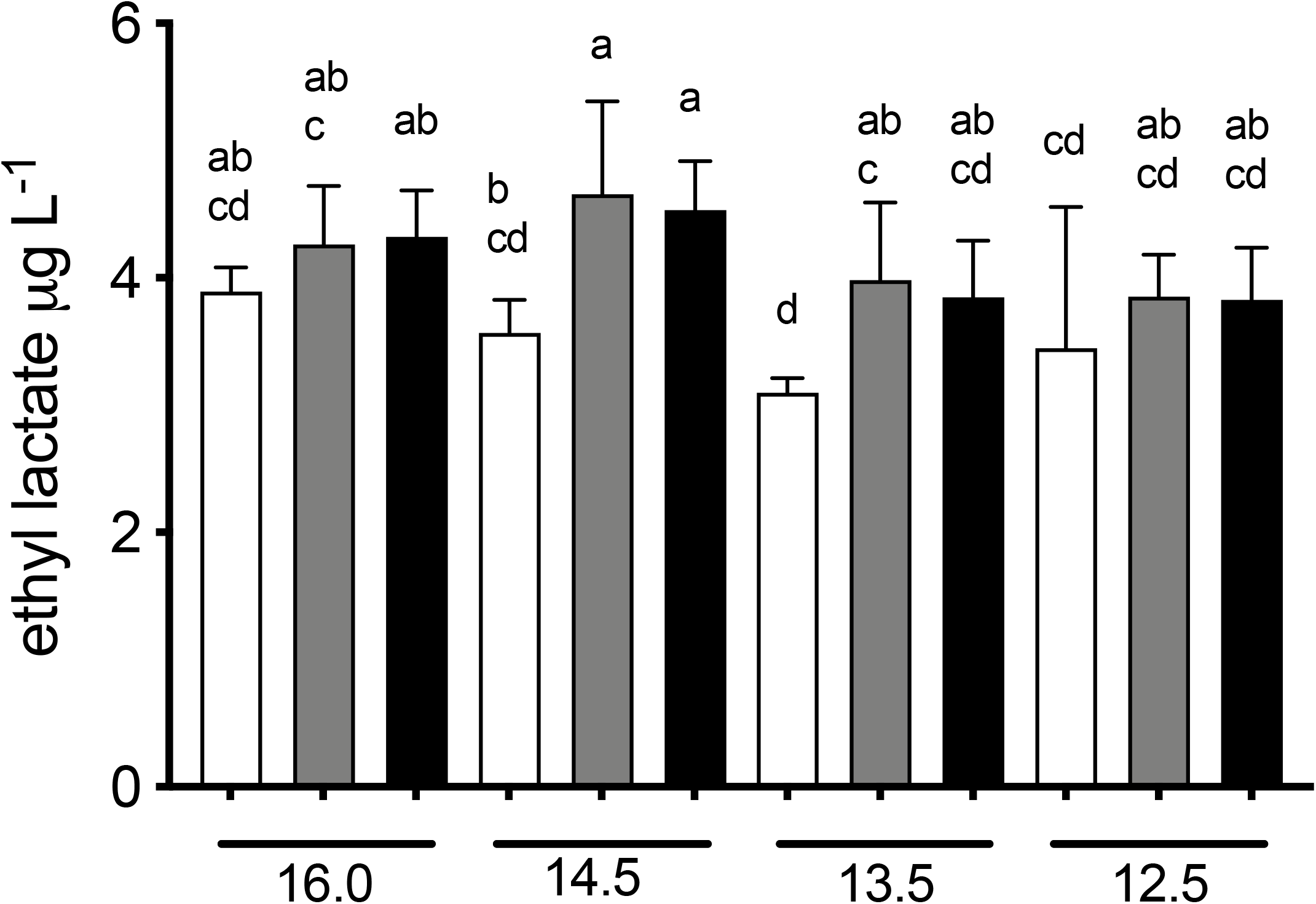
Ethyl lactate measured in wines (GC-MS) fermented by Lalvin R2 ™ with no malolactic bacteria added. Wines were made from juice with initial Bé of 16 or diluted with water to 14.5, 13.5 or 12.5. Nutrient was also added to juices; 0 (□), 200 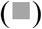 or 400 (■) mg L^−1^. Dashed line represents aroma threshold of isoamyl acetate (30 μg L^−1^ (Guth, 1997)). Significantly different values are labelled with different letters or letter groups (a-d), one way ANOVA, p < 0.05).

## Notes

### Competing Interest Statement

The authors have declared no competing interest.

## References

Bell, S. J., & Henschke, P. A. (2005). Implications of nitrogen nutrition for grapes, fermentation and wine. Australian Journal of Grape and Wine Research, 11(3), 242–295. https://doi.org/10.1111/j.1755-0238.2005.tb00028.x.

Bloem, A., Sanchez, I., Dequin, S., & Camarasa, C. (2016). Metabolic impact of redox cofactor perturbations on the formation of aroma compounds in *Saccharomyces cerevisiae*. Appl Environ Microbiol, 82(1), 174–183. https://doi.org/10.1128/AEM.02429-15.

Boss, P. K., Pearce, A. D., Zhao, Y. J., Nicholson, E. L., Dennis, E. G., & Jeffery, D. W. (2015). Potential grape-derived contributions to volatile ester concentrations in wine. Molecules, 20(5), 7845–7873. https://doi.org/10.3390/molecules20057845.

Chen, Y., Siewers, V., & Nielsen, J. (2012). Profiling of cytosolic and peroxisomal Acetyl-CoA metabolism in *Saccharomyces cerevisiae*. PLoS One, 7(8), e42475. https://doi.org/10.1371/journal.pone.0042475.

Cowey, G. (2017). Making water additions to high sugar must. AWRI Technical Review, 227, 9–12.

Duffour, J., Malcorps, P., & Silcock, P. (2003). Control of Ester Synthesis During Brewery Fermentation. In Brewing Yeast Fermentation Performance (pp. 213–233).

Escudero, A., Campo, E., Fariña, L., Cacho, J., & Ferreira, V. (2007). Analytical characterization of the aroma of five premium red wines. insights into the role of odor families and the concept of fruitiness of wines. Journal of Agricultural and Food Chemistry, 55(11), 4501–4510. https://doi.org/10.1021/jf0636418.

Etievant, P. X. (1991). Volatile compounds in foods and beverages. In H. Maarse (Ed.), Food science and technology; 44 (pp. 483–546). New York, N.Y: Marcel Dekker.

Ferreira, V., Lopez, R., & Cacho, J. F. (2000). Quantitative determination of the odorants of young red wines from different grape varieties. Journal of the Science of Food and Agriculture, 50(11), 1659–1667. https://doi.org/Doi10.1002/1097-0010(20000901)80:11<1659::Aid-Jsfa693>3.0.Co;2-6.

Gomez-Miguez, M. J., Cacho, J. F., Ferreira, V., Vicario, I. M., & Heredia, F. J. (2007). Volatile components of Zalema white wines. Food Chemistry, 100(4), 1464–1473. https://doi.org/10.1016/j.foodchem.2005.11.045.

Guth, H. (1997). Quantitation and sensory studies of character impact odorants of different white wine varieties. Journal of Agricultural and Food Chemistry, 45(8), 3027–3032. https://doi.org/DOI10.1021/jf970280a.

Harbertson, J. F., Mireles, M. S., Harwood, E. D., Weller, K. M., & Ross, C. F. (2009). Chemical and sensory effects of saignée, water addition, and extended maceration on high brix must. American Journal of Enology and Viticulture, 60(4), 450–460. https://www.ajevonline.org/content/ajev/60/4/450.full.pdf.

Hazelwood, L. A., Daran, J. M., van Maris, A. J., Pronk, J. T., & Dickinson, J. R. (2008). The Ehrlich pathway for fusel alcohol production: a century of research on *Saccharomyces cerevisiae* metabolism. Appl Environ Microbiol, 74(8), 2259–2266. http://www.ncbi.nlm.nih.gov/entrez/query.fcgi?cmd=Retrieve&db=PubMed&dopt=Citation&list_uids=18281432.

Hernández-Orte, P., Ibarz, M. J., Cacho, J., & Ferreira, V. (2005). Effect of the addition of ammonium and amino acids to musts of Airen variety on aromatic composition and sensory properties of the obtained wine. Food Chemistry, 89(2), 163–174. https://doi.org/10.1016/j.foodchem.2004.02.021.

Hranilovic, A., Li, S., Boss, P. K., Bindon, K., Ristic, R., Grbin, P. R., Van der Westhuizen T., & Jiranek, V. (2018). Chemical and sensory profiling of Shiraz wines co-fermented with commercial *non-Saccharomyces* inocula. Australian Journal of Grape and Wine Research, 24(2), 166–180. https://doi.org/10.1111/ajgw.12320.

Jiang, J., Sumby, K. M., Sundstrom, J. F., Grbin, P. R., & Jiranek, V. (2018). Directed evolution of *Oenococcus oen*i strains for more efficient malolactic fermentation in a multi-stressor wine environment. Food Microbiol, 73, 150–159. https://doi.org/10.1016/j.fm.2018.01.005.

Lallemand. (2019). Fermaid O Natural Yeast Derived Nutrient. Retrieved from: https://catalogapp.lallemandwine.com/uploads/nutrient/docs/f4a473f989e161ecc34cae1cedd6058cefb223b2.pdf. Accessed 1.7.20.

Lin, M. M.-H., Boss, P. K., Walker, M. E., Sumby, K. M., Grbin, P. R., & Jiranek, V. (2020). Evaluation of indigenous *non-Saccharomyces* yeasts isolated from a South Australian vineyard for their potential as wine starter cultures. International Journal of Food Microbiology, 312, 108373. https://doi.org/10.1016/j.ijfoodmicro.2019.108373.

Moreno, J. A., Zea, L., Moyano, L., & Medina, M. (2005). Aroma compounds as markers of the changes in sherry wines subjected to biological ageing. Food Control, 16(4), 333–338. https://doi.org/10.1016/j.foodcont.2004.03.013.

Peinado, R. A., Moreno, J., Bueno, J. E., Moreno, J. A., & Mauricio, J. C. (2004). Comparative study of aromatic compounds in two young white wines subjected to pre-fermentative cryomaceration. Food Chemistry, 84(4), 585–590. https://doi.org/10.1016/S0308-8146(03)00282-6.

Petrie, P. R., Teng, B., Smith, P. A., & Bindon, K. (2019). Managing high Baume juice using dilution. Wine and Viticulture Journal, Summer, 36–37.

Pozo-Bayón, M. Á., Andújar-Ortiz, I., & Moreno-Arribas, M. V. (2009). Volatile profile and potential of inactive dry yeast-based winemaking additives to modify the volatile composition of wines. Journal of the Science of Food and Agriculture, 89(10), 1665–1673. https://doi.org/10.1002/jsfa.3638.

Pronk, J. T., Yde Steenema, H., & Van Dijken, J. P. (1996). Pyruvate Metabolism in *Saccharomyces cerevisiae*. Yeast, 12(16), 1607–1633. https://doi.org/10.1002/(sici)1097-0061(199612)12:16<1607::Aid-yea70>3.0.Co;2-4.

Remize, F., Andrieu, E., & Dequin, S. (2000). Engineering of the pyruvate dehydrogenase bypass in *Saccharomyces cerevisiae*: role of the cytosolic Mg(^2+^) and mitochondrial K(^+^) acetaldehyde dehydrogenases Ald6p and Ald4p in acetate formation during alcoholic fermentation. Applied and Environmental Microbiology, 66(8), 3151–3159. https://doi.org/10.1128/aem.66.8.3151-3159.2000.

Saerens, S. M. G., Delvaux, F., Verstrepen, K. J., Van Dijck, P., Thevelein, J. M., & Delvaux, F. R. (2008). Parameters affecting ethyl ester production by *Saccharomyces cerevisiae* during fermentation. Applied and Environmental Microbiology, 74(2), 454–461. https://doi.org/10.1128/aem.01616-07.

Saerens, S. M. G., Verstrepen, K. J., Van Laere, S. D. M., Voet, A. R. D., Van Dijck, P., Delvaux, F. R., & Thevelein, J. M. (2006). The *Saccharomyces cerevisiae EHT1* and *EEB1* genes encode novel enzymes with medium-chain fatty acid ethyl ester synthesis and hydrolysis capacity. Journal of Biological Chemistry, 281(7), 4446–4456. https://doi.org/10.1074/jbc.M512028200.

Salo, P. (1970). Determining the odor thresholds for some compounds in alcoholic beverages. Journal of Food Science, 35(1), 95–99. https://doi.org/10.1111/j.1365-2621.1970.tb12378.x.

Schelezki, O. J., Antalick, G., Šuklje, K., & Jeffery, D. W. (2020). Pre-fermentation approaches to producing lower alcohol wines from Cabernet Sauvignon and Shiraz: Implications for wine quality based on chemical and sensory analysis. Food Chemistry, 309, 125698. https://doi.org/10.1016/j.foodchem.2019.125698.

Schelezki, O. J., Smith, P. A., Hranilovic, A., Bindon, K. A., & Jeffery, D. W. (2018). Comparison of consecutive harvests versus blending treatments to produce lower alcohol wines from Cabernet Sauvignon grapes: Impact on polysaccharide and tannin content and composition. Food Chemistry, 244, 50–59. https://doi.org/10.1016/j.foodchem.2017.10.024.

Schelezki, O. J., Suklje, K., Boss, P. K., & Jeffery, D. W. (2018). Comparison of consecutive harvests versus blending treatments to produce lower alcohol wines from Cabernet Sauvignon grapes: Impact on wine volatile composition and sensory properties. Food Chemistry, 259, 196–206. https://doi.org/10.1016/j.foodchem.2018.03.118.

Schultz, H. R. (2016). Global climate change, sustainability, and some challenges for grape and wine production. Journal of Wine Economics, 11(1), 181–200. https://doi.org/10.1017/jwe.2015.31.

Takeoka, G., Buttery, R. G., Flath, R. A., Teranishi, R., Wheeler, E. L., Wieczorek, R. L., & Guentert, M. (1989). Volatile Constituents of Pineapple *Ananas Comosus* [L.] Merr.). In Flavor Chemistry (pp. 223–237): American Chemical Society.

Taylor, G. T., & Kirsop, B. H. (1977). The origin of the medium chain length fatty acids in beer. Journal of the Institute of Brewing, 83(4), 241–243. https://doi.org/10.1002/j.2050-0416.1977.tb03802.x.

Teng, B., Petrie, P. R., Smith, P. A., & Bindon, K. A. (2020). Comparison of water addition and early-harvest strategies to decrease alcohol concentration in *Vitis vinifera* cv. Shiraz wine: impact on wine phenolics, tannin composition and colour properties. Australian Journal of Grape and Wine Research, 26(2), 158–171. https://doi.org/10.1111/ajgw.12430.

Torrea, D., Fraile, P., Garde, T., & Ancin, C. (2003). Production of volatile compounds in the fermentation of chardonnay musts inoculated with two strains of *Saccharomyces cerevisiae* with different nitrogen demands. Food Control, 14(8), 565–571. http://www.sciencedirect.com/science/article/B6T6S-482G9D2-D/2/d6c8022a366b98e79e0e669480a62401.

Torrea, D., Varela, C., Ugliano, M., Ancin-Azpilicueta, C., Leigh Francis, I., & Henschke, P. A. (2011). Comparison of inorganic and organic nitrogen supplementation of grape juice – Effect on volatile composition and aroma profile of a Chardonnay wine fermented with *Saccharomyces cerevisiae* yeast. Food Chemistry, 127(3), 1072–1083. https://doi.org/10.1016/j.foodchem.2011.01.092.

Varela, J. C. S., & Mager, W. H. (1996). Response of *Saccharomyces cerevisiae* to changes in external osmolarity. Microbiology, 142(4), 721–731. https://doi.org/10.1099/00221287-142-4-721.

Walker, M. E., Nguyen, T. D., Liccioli, T., Schmid, F., Kalatzis, N., Sundstrom, J. F., Gardner J. M., & Jiranek, V. (2014). Genome-wide identification of the Fermentome; genes required for successful and timely completion of wine-like fermentation by *Saccharomyces cerevisiae*. BMC Genomics, 15, 17. https://doi.org/10.1186/1471-2164-15-552.

Xu, X., Bao, M., Niu, C., Wang, J., Liu, C., Zheng, F., Yongxian, L., & Li, Q. (2019). Engineering the cytosolic NADH availability in lager yeast to improve the aroma profile of beer. Biotechnology Letters, 41(3), 363–369. https://doi.org/10.1007/s10529-019-02653-x.

Zhang, K., Sawaya, M. R., Eisenberg, D. S., & Liao, J. C. (2008). Expanding metabolism for biosynthesis of nonnatural alcohols. Proceedings of the National Academy of Sciences, 105(52), 20653–20658. https://doi.org/10.1073/pnas.0807157106.

